# Mechanical confinement induces ferroptosis through mitochondrial dysfunction

**DOI:** 10.1101/2024.02.23.581510

**Authors:** Fang Zhou, Robert J. Ju, Chenlu Kang, Jiayi Li, Ao Yang, Alexandre Libert, Yujie Sun, Ling Liang, Xiaoqing Hu, Samantha J. Stehbens, Congying Wu

**Affiliations:** Institute of Systems Biomedicine, School of Basic Medical Sciences, Peking University Health Science Center, Beijing, 100191 China; The University of Queensland, Institute for Molecular Bioscience, St Lucia Brisbane, Queensland, Australia; The University of Queensland, Australian Institute for Bioengineering and Nanotechnology, St Lucia Brisbane, Queensland, Australia; Department of Biochemistry and Biophysics, School of Basic Medical Sciences, Peking University Health Science Center, Beijing, 100191, China; State Key Laboratory of Membrane Biology, Biomedical Pioneer Innovation Center (BIOPIC), School of Life Sciences, Peking University, Beijing, 100871, China; Department of Sports Medicine, Institute of Sports Medicine of Peking University, Beijing Key Laboratory of Sports Injuries, Peking University Third Hospital, Beijing, 100191, China; International Cancer Institute, Peking University, Beijing, 100191, China

## Abstract

Cells exist in highly crowded environments where they are exposed to fluctuating mechanical forces arising from surrounding cells and the extracellular matrix microenvironment. In these settings, external forces are transmitted to intracellular organelles including the nucleus. While cells can survive confinement, extended duration of confinement or confinement in settings where cells are unable to escape, can result in cell death. How cells sense and respond to prolonged confinement to trigger cell death remains unclear. Here, we demonstrate that nuclear deformation generated by axial confinement triggers the programmed cell death pathway – ferroptosis. We show that axial confinement results in Drp1-dependent mitochondrial fragmentation and cPLA2 translocation to mitochondria, where Drp1 undergoes acute phase separation. Ensuing mitochondrial ROS accumulation and arachidonic acid production concertedly leads to lipid peroxidation evoking ferroptosis. Finally, of clinical relevance, we find that in human osteoarthritis tissue cPLA2 exhibits mitochondrial localization and high ROS levels. Together, our findings unveil a pivotal role for Drp1 and cPLA2 in linking mechanical confinement with mitochondrial dysfunction resulting in ferroptosis, which sheds new light on a mechanical mechanism of pathophysiology in osteoarthritis.

## Introduction

In multicellular organisms, cells are exposed to mechanical forces including shear stress and compression^1, 2^. In the context of confined migration, the squeezing of cells through narrow constrictions results in high-levels of nuclear deformation triggering mesenchymal-to-amoeboid migration mode switching^3, 4^. However, not all cells are motile and how adherent cells respond, adapt to, and survive persistent mechanical force at the molecular level, is less clear.

Unlike chemical ligands that rely on cell surface receptors to transduce signals into the cell, mechanical force can propagate throughout cells via deformation of intracellular structures^5, 6^. Within cells, the cytoskeleton and organelles are mechano-responsive, acting in combination with molecular-scale alterations in force sensitive proteins to modify cellular responses to mechanical force^7, 8^. The role of the nucleus as a mechanoresponsive organelle is the most well described, with external force resulting in transcription factor nuclear translocation^9^, and changes in chromatin rheology and architecture to protect genomic content^10^. Recently, nuclear deformation was shown to activate contractility of the actomyosin cortex, facilitating the modulation of cell migration in response to force^11, 12^. Further, studies have found when cells migrate through pores that distend and herniate the nuclear membrane can result in nuclear rupture, DNA damage and be linked to carcinogenic and inflammogenic status^13–15^. Immune cells can detect inflammatory signals released via the programmed forms of immunogenic cell deaths - pyroptosis, necroptosis, and ferroptosis^16, 17^. Interestingly, ferroptosis has recently been observed in osteoarthritis (OA), a disease intimately associated with mechanical overloading^18^. Thus, we hypothesised that chronic inflammation associated with OA could be a result of ferroptosis within mechanically overloaded cells experiencing prolonged confinement.

To investigate this we used an in-vitro model of axial confinement and found that nuclear deformation via axial confinement triggers ferroptosis. We observed cPLA2 translocation to mitochondria followed by arachidonic acid (ARA) generation, and enhanced Drp1 phase separation followed by mitochondrial fragmentation and accumulation of mitochondrial ROS (mtROS). These concertedly lead to lipid oxidation and ferroptosis-mediated cell death. Finally, we have demonstrated that OA patient samples show characteristic of this mechano-stress response.

## Results

### 1. Mechanical confinement triggers ferroptosis

To first understand how axial confinement affects cell morphology and survival, we performed dynamic axial confinement of HeLa cells to 3 µm height using microfabricated PolyDiMethylSiloxane (PDMS) micropillars bonded to glass coverslips^19^. At this height, the nucleus is sufficiently deformed to activate cortical actomyosin contractility^3, 11^. When cells were subjected to continuous axial confinement for a duration of 9 hours, cells began to rupture and die at 3 hours, with continuous increases in cell death persisting through to 9 hours, as indicated by nuclear propidium iodide (PI) influx (Figure 1A-1B, Movie S1). In the absence of confinement height control (beneath the PDMS micropillars), cells instantly ruptured exhibiting acute influx of nuclear PI signal (Figure S1A). In addition, we observed the same phenomenon when using a second method of axial confinement^20^ using 3 μm microspheres and a machined stainless steel weight to confine cells (Figure S1B). Cells could also viably sustain several hours of axial confinement, exhibiting a distended morphology marked by large plasma membrane blebs, which appeared and resolved, until cell rupture was detected at 6 hours confinement, where above 80% of the cells were PI positive (Figure S1C-S1D). This suggested that cells exposed to controlled ∼3 µm axial confinement underwent a slower, likely programmed, process of cell death.

**Figure 1.**
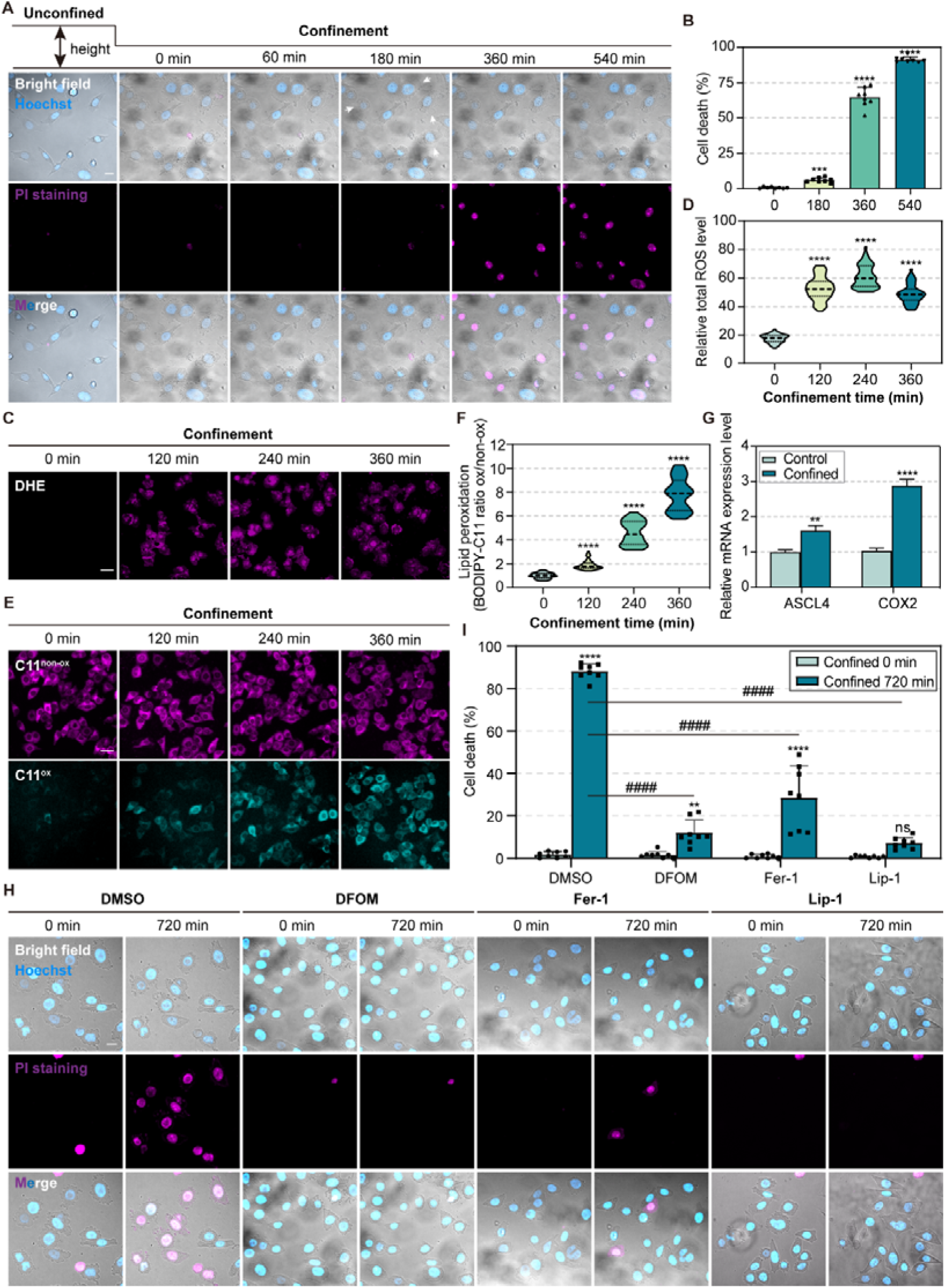
Mechanical confinement triggers ferroptosis. (A) Representative time-lapse images of HeLa cells during confinement in the presence of propidium iodide (PI) (magenta, 10 μM). Nuclei labelled with Hoechst (blue, 5 μg/mL for 20 min). White arrow: large plasma membrane blebs. Scale bar = 20 μm. (B) Quantification of the percentage of cell death in HeLa cells during confinement (*N* = 8 fields, *\*\*\*p* < 0.001, *\*\*\*\*p* < 0.0001). The cell death is measured by the PI positive nuclei/total nuclei * 100. (C) Representative time-lapse images of dihydroethidium (DHE) staining (magenta, 10 μM for 1 hour) indicated total ROS level in HeLa cells during confinement. Scale bar = 40 μm. (D) Quantification of total ROS level in HeLa cells during confinement (*N* = 50 cells, *\*\*\*\*p* < 0.0001). Violin plot center lines denote the median, and dashed lines represent the 25th and 75th percentiles. (E) Representative time-lapse images showing BODIPY-C11 581/591 staining (5 μM for 1 hour) indicated lipid ROS level in HeLa cells during confinement. Non-oxidative state (magenta), oxidative state (cyan). Scale bar = 40 μm. (F) Quantification of lipid ROS level in HeLa cells during confinement (*N* = 50 cells, *\*\*\*\*p* < 0.0001). Data was calculated by BODIPY-C11 fluorescence intensity ratio of oxidative/non-oxidative. (G) qRT-PCR assay showing the ASCL4 and COX2 mRNA levels in HeLa cells upon confined and unconfined conditions (*\*\*p* < 0.01, *\*\*\*\*p* < 0.0001). (H) Representative time-lapse images of HeLa cells treated with DMSO or ferroptosis inhibitors during 3 μm confinement mediated by PDMS micropillars in the presence of PI, including DFOM (100 μM), Ferrostatin-1 (Fer-1, 1 μM), Liproxstatin-1 (Lip-1, 0.5 μM). Scale bar = 20 μm. (I) The percentage of cell death in HeLa cells treated with DMSO or ferroptosis inhibitors during confinement (*N* = 8 fields, ns represents not significant, *\*\*p* < 0.01, *\*\*\*\*p* < 0.0001, *^####^p* < 0.0001).

To exclude that this cell death was not caused by hypoxia during confinement, we performed confinement using 20 μm microsphere spacers where cells were not in direct contact with the confinement surface^19^. Under 20 μm axial confinement height, no observable cell death was detected (Figure S1E), indicating that the cell death we observed was in response to prolonged confinement conditions and not hypoxia induced.

We postulated that the large membrane distention we observed during axial confinement could be in line with the characteristic membrane blebbing during the programmed cell death pathway, pyroptosis^21, 22^. However, when we screened for activation of the classical effectors of pyroptosis, we failed to detect the cleavage of the pore-forming effectors, GSDMD or GSDME (Figure S1F). Moreover, the pan-caspase inhibitor Z-VAD-FMK used to stall pyroptosis^23, 24^, failed to block this process (Figure S1G). In addition, we detected no changes in MLKL phosphorylation^25^, eliminating the possibility of confinement induced necroptosis (Figure S1H). In line with this, NSA-dependent inhibition of necroptosis failed to block confinement-induced cell death (Figure S1I). Interestingly, alongside PI influx of confined cells, we also observed a time-dependent increase in total and lipid ROS using the fluorescent probes DHE and BODIPY-C11, respectively. Total and lipid ROS were observed to accumulate at 2 hours under 3 μm (Figure 1C-1F), but not 20 μm confinement (Figure S1J). As lipid peroxidation is a hallmark of ferroptosis^26, 27^, we examined additional markers of activation of the ferroptotic pathway. We observed that RNA expression of the ferroptosis biomarkers, acyl-CoA synthetase long-chain family member 4 (ACSL4) and cyclooxygenase-2 (COX2) increased after confinement (Figure 1G). Remarkably, we found that confinement induced-cell death was significantly diminished when cells were confined in the presence of ferroptosis inhibitors, including DFOM, Ferrostatin-1 (Fer-1), Liproxstatin-1 (Lip-1), and Trolox, (Figure 1H-1I, Figure S1K, Movie S2). Together our data demonstrate that cells subjected to prolonged axial confinement exhibit programmed cell death via ferroptosis.

### 2. cPLA2 is activated and translocates to mitochondria upon confinement

To understand how axial confinement molecularly triggers ferroptosis activation, we next asked whether mechanically responsive cell surface proteins might play a role in this response. Mechano-sensitive ion channels^28^ and cell-adhesion molecules^8^ constitute two canonical mechanosensors in cells. When we performed inhibition of Piezo with GsMTx4 and depletion of integrins with siRNAs – we observed no changes in cell death (Figure S2A-S2B). We next examined the nucleus which acts as a cellular ruler^11, 12^, capable of converting structural changes via mechanical input into signaling or transcriptional output^29^. We confirmed nuclear deformation occurred with 3 μm (cf. 20 μm) confinement, observing a 35% increase in the nuclear projected surface area (Figure 2A-2B).

**Figure 2.**
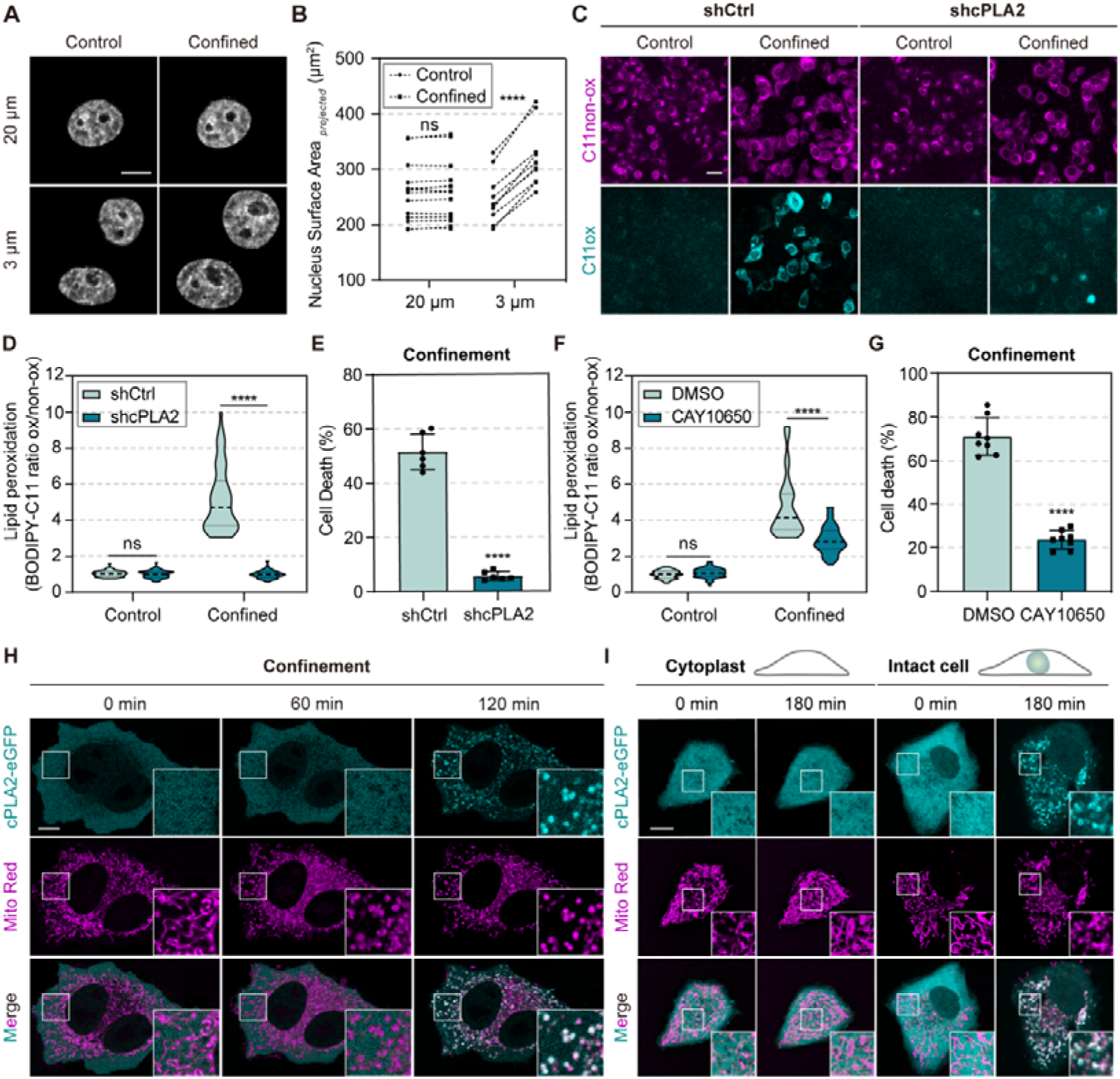
cPLA2 is activated and translocates to mitochondria upon confinement. (A) xy views of the Hoechst-stained nucleus before and after 20 μm (top) or 3 μm (bottom) confinement. Scale bar = 10 μm. (B) Quantification of projected nucleus surface area upon confinement (*N*_20_ μ_m_ = 14 nuclei, *N*_3_ μ_m_ = 10 nuclei, ns represents not significant, *\*\*\*\*p* < 0.0001). (C) Representative images of BODIPY-C11 581/591 staining in control (shCtrl) and cPLA2 knockdown (shcPLA2) HeLa cells in the absence (control) and presence (confined) of confinement. Scale bar = 40 μm. (D) Quantification of lipid ROS level in shCtrl and shcPLA2 HeLa cells (*N* = 50 cells, ns represents not significant, *\*\*\*\*p* < 0.0001). (E) The percentage of cell death in shCtrl and shcPLA2 HeLa cells upon confinement (*N* = 6 fields, *\*\*\*\*p* < 0.0001). (F) Lipid ROS levels in HeLa cells treated with DMSO (control) or CAY10650 (150 nM, cPLA2 inhibitor) (*N*_DMSO_ = 60 cells, *N*_CAY10650_ = 66 cells, ns represents not significant, *\*\*\*\*p* < 0.0001). (G) The percentage of cell death in HeLa cells treated with DMSO or CAY10650 upon confinement (*N* = 8 fields, *\*\*\*\*p* < 0.0001). (H) Representative time-lapse images of cPLA2-eGFP expressing HeLa cells co-stained with MitoTrackerTM Red CMXRos (Mito Red, 200 nM) for 30 min during confinement. Scale bar = 10 μm. (I) Representative images of cPLA2-eGFP and mitochondria (Mito Red) spatio-temporal localization in intact HeLa cells or enucleated cytoplasts upon confinement. Scale bar = 10 μm.

Nuclear deformation has been shown to induce translocation of the activate cytosolic phospholipase 2 (cPLA2) to the nuclear envelope^30^, where activated cPLA2 catalyzes production of ARA^11, 12^. Importantly, during this process lipid hydroperoxides accumulate, which are prerequisites for ferroptosis^31^. Separately, it is reported that cPLA2 is capable of localizing to mitochondria for activation following sphingolipid lactosylceramide treatment in astrocytes^32^. Thus, we asked whether cPLA2 may play a role in confinement-induced ferroptosis. When we depleted cells of cPLA2 using shRNA-mediated knock-down (Figure S2C), we observed a reduction in lipid ROS level and abrogation of cell death in confined cells (Figure 2C-2E). Similarly, we could reproduce this result with the cPLA2 inhibitor CAY10650 (Figure 2F-2G), supporting the catalytic role of cPLA2 in confinement-induced ferroptosis. Strikingly, when we performed live-cell imaging of cPLA2-eGFP dynamics in cells subjected to axial confinement, we did not observe nuclear envelope translocation (Figure S2D). Instead, we observed cPLA2 translocation to mitochondria beginning at 90 min of 3 μm axial confinement, but not 20 μm (Figure 2H, Figure S2E, Movie S3). To demonstrate that confinement triggers cPLA2 translocation from the cytoplasm to the mitochondria, we used centrifugation-based enucleation^33, 34^ of cells to produce cytoplasts (Figure S2F). In cytoplasts exposed to axial confinement, cPLA2 translocation to the mitochondria was abolished (Figure 2I, Figure S2G). These data suggest that nuclear deformation triggers cPLA2 translocation to mitochondria, leading to an increase in lipid ROS consistent with activation of ferroptosis.

### 3. Confinement elevates mitochondrial fission and mtROS

While observing cPLA2 mitochondrial translocation, we noted that the majority of cells (87%) exhibited mitochondrial fragmentation when exposed to 3 μm confinement (Figure S3A-S3B). We thus performed live-cell time lapse imaging of mitochondria labeled cells (Tom20-mCherry) under axial confinement, and observed mitochondrial swelling and fragmentation (Figure 3A-3B, Movie S4). We also detected mitochondrial membrane depolarization during confinement (Figure 3C-3D), using tetramethylrhodamine methyl (TMRM). Importantly, as we did not observe Parkin translocation to mitochondria (Figure S3C), we were able to exclude mitophagy. As mitochondria defects are responsible for increased mtROS^35^, which has been implicated in lipid peroxidation and ferroptosis onset^36^, we next examined whether mtROS levels were also increased during confinement-induced mitochondrial fission. Using Mito-roGFP2, we observed a significant increase of mtROS when cells underwent axial confinement (Figure 3E-3F). When we inhibited mtROS accumulation (Figure S3D) by using MitoTEMPO, a mitochondria-targeted superoxide dismutase mimetic, we found significantly reduced total ROS and lipid ROS, as well as decreased cell death of confined cells (Figure 3G-3I). We observed a similar effect with the antioxidant EUK134 or NAC (Figure S3E-S3F), suggesting that mtROS plays a key role in modulating confinement-induced ferroptosis.

**Figure 3.**
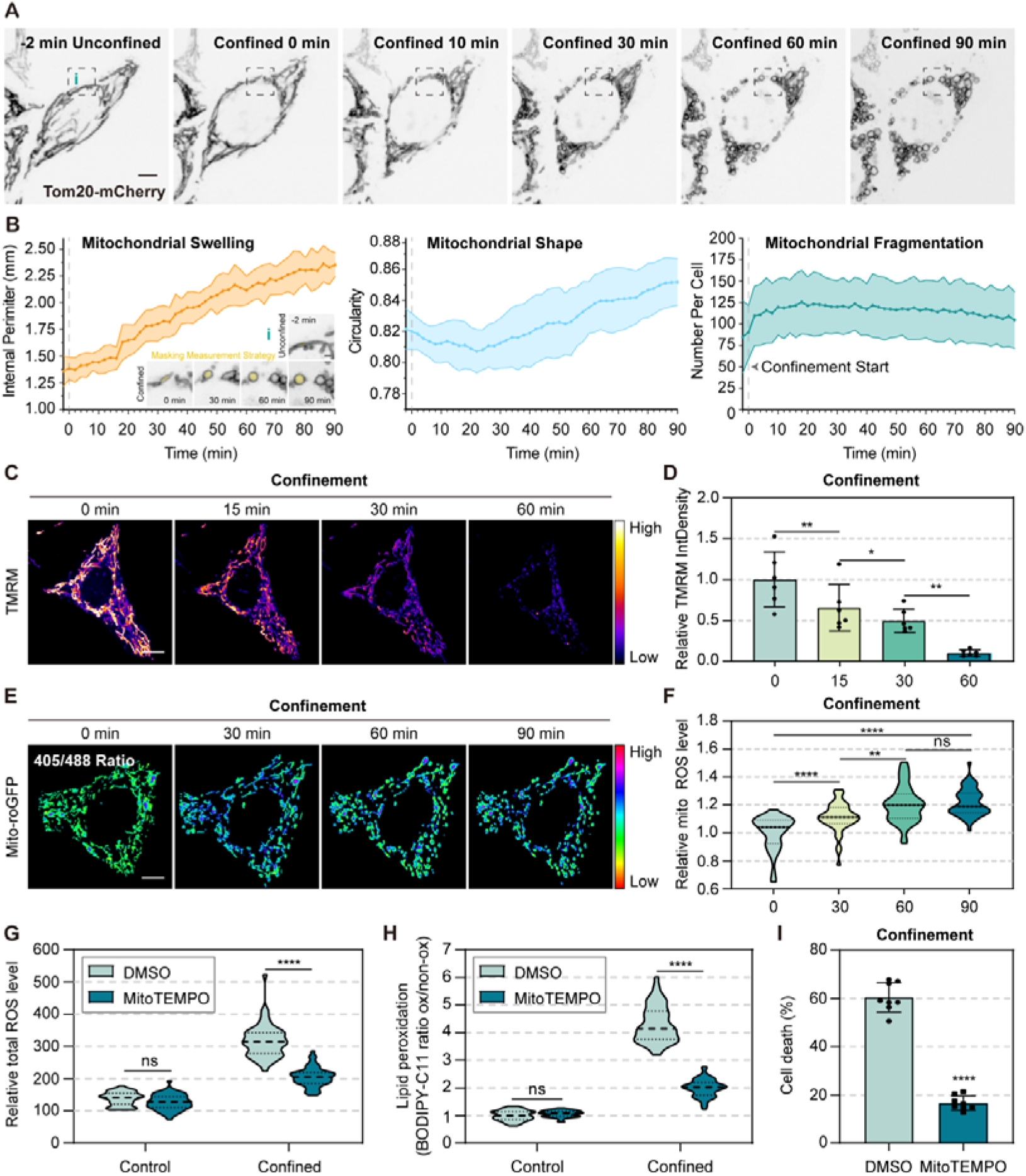
Mitochondrial fragmentation and elevated mtROS level contribute to confinement-induced ferroptosis. (A) Representative time-lapse images of mitochondria in HeLa cells transfected with Tom20-mCherry under 3 μm high micropillars. Scale bar = 10 μm. (B) Quantification of mitochondrial morphology in (A) at 3 μm confinement for 90 min (*N* = 12 cells). Magnified images are areas outlined (i), black dashed boxes of (A). (C) Representative time-lapse images of TMRM staining (100 nM, 20 min) in HeLa cells during 60 min confinement, fluorescence intensity presented as a heatmap (low-high). Scare bar = 10_μm. (D) Quantification of TMRM fluorescence intensity per cell under confinement, normalized by data at 0 min (*N* = 6 cells, *\*p* < 0.05, *\*\*p* < 0.01). (E) Representative time-lapse ratiometric images showing mtROS level in HeLa cells stably expressing Mito-roGFP during 90 min confinement. Scare bar = 10_μm. (F) Quantification of the relative mtROS level per cell in HeLa cells under confinement, normalized by data at 0 min (*N* = 44 cells, ns represents not significant, *\*\*p* < 0.01, *\*\*\*\*p* < 0.0001). (G)Total ROS level in HeLa cells treated with DMSO or MitoTEMPO (10 μm) in confined and unconfined conditions (*N* = 50 cells, ns represents not significant, *\*\*\*\*p* < 0.0001). (H) Lipid ROS level in HeLa cells treated with DMSO or MitoTEMPO in confined and unconfined conditions (*N* = 50 cells, ns represents not significant, *\*\*\*\*p* < 0.0001). (I) The percentage of cell death in HeLa cells treated with DMSO or MitoTEMPO upon confinement (*N* = 8 fields, *\*\*\*\*p* < 0.0001).

Our observation of confinement-induced mitochondrial fission recapitulates reports of altered mitochondrial morphology in ferroptotic cells^26, 37^. To better understand the temporal dynamics of mitochondrial fission in response to nuclear compression, we performed atomic force microscopy (AFM) with a flat tipless cantilever to deform single cells above the nucleus, while monitoring mitochondrial morphology by live-cell imaging. Notably, we observed rapid mitochondrial fission when the cantilever deformed the nucleus (Figure 4A). Fission was absent in mitochondria-containing cytoplasts (Figure S4A), in line with our earlier observations of compression-induced fission being a nuclear-dependent process (Figure 4B).

**Figure 4.**
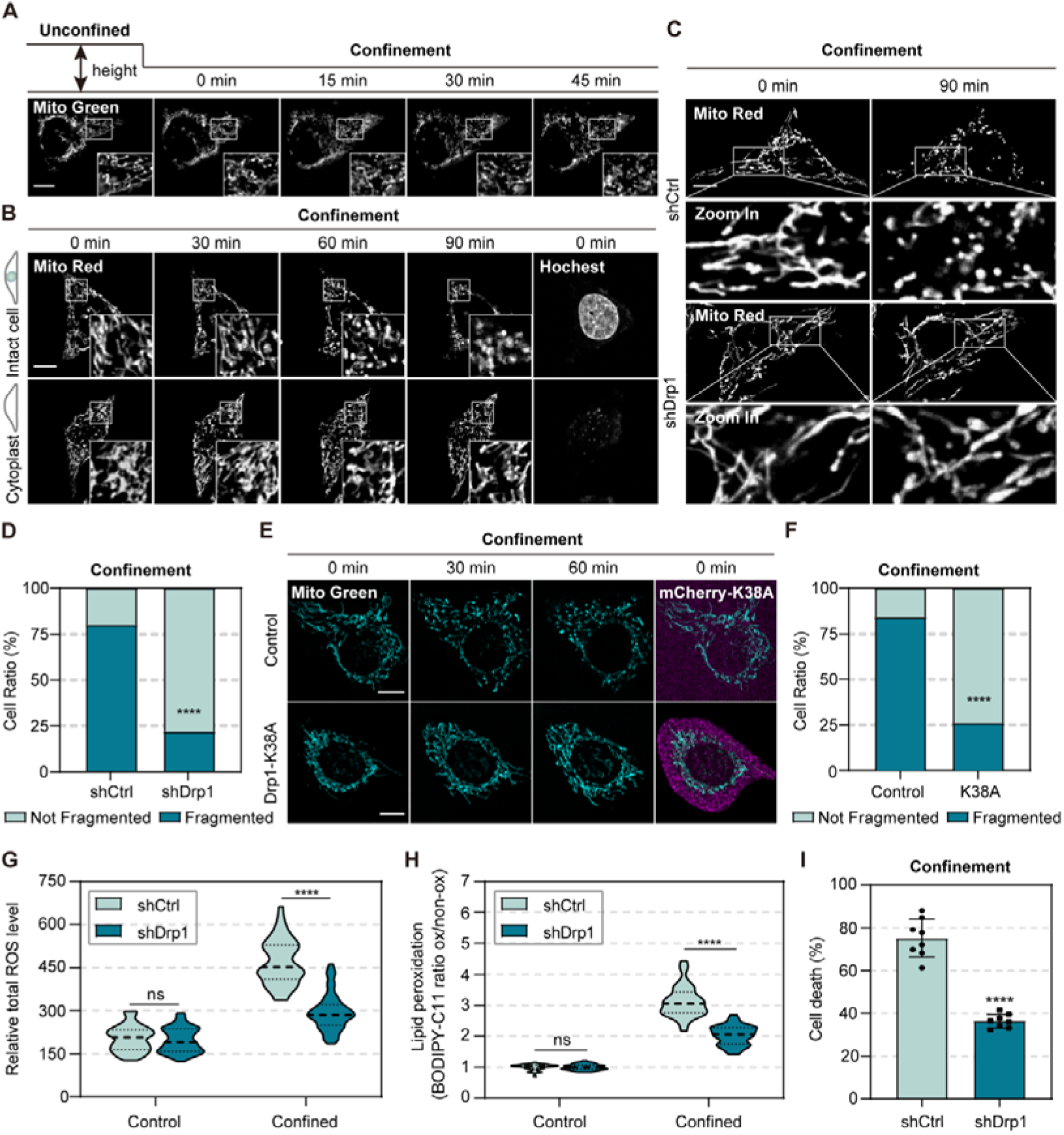
Nuclear confinement-triggered mitochondrial fragmentation is dependent on Drp1. (A) Representative time-lapse images of mitochondria (Mito Green) in a HeLa cell compressed by AFM. Insets are magnified images of areas indicated by white boxes. Scale bar = 10 μm. (B) Representative time-lapse images of mitochondria (Mito Red) in intact HeLa cells or enucleated cytoplasts during 90 min confinement. Insets are magnified images of areas indicated by white boxes. Scale bar = 10 μm. (C) Representative images of mitochondria (Mito Red) in control (shCtrl) and Drp1 knock-down (shDrp1) HeLa cells after confinement for 0 or 90 min. Insets are magnified images of areas indicated by white boxes. Scale bar = 10 μm. (D) The percentage of cells with fragmented mitochondria in shCtrl and shDrp1 HeLa cells upon 90 min confinement (*N*_shCtrl_ = 56 cells, *N*_shDrp1_ = 49 cells, *\*\*\*\*p* < 0.0001, Fisher’s exact test). (E) Representative time-lapse images of mitochondria (Mito Green) in HeLa cells expressing mCherry-Drp1-K38A during 60 min confinement. Non-transfected HeLa cells in the same sample were used as a control. Scale bar = 10 μm. (F) The percentage of cells with fragmented mitochondria in HeLa cells expressing mcherry-Drp1-K38A or in the control group upon 60 min confinement (*N* = 50 cells, *\*\*\*\*p* < 0.0001). (G) Total ROS level in shCtrl and shDrp1 HeLa cells upon confined and unconfined conditions (*N* = 50 cells, ns represents not significant, *\*\*\*\*p* < 0.0001). (H) Lipid ROS level in shCtrl and shDrp1 HeLa cells upon confined and unconfined conditions (*N* = 40 cells, ns represents not significant, *\*\*\*\*p* < 0.0001). (I) The percentage of cell death in shCtrl and shDrp1 HeLa cells upon confinement (*N* = 8 cells, *\*\*\*\*p* < 0.0001).

Canonically, mitochondrial fission is executed by the dynamin-like GTPase Drp1^38^. To assess whether Drp1 was involved in confinement-mediated mitochondrial fission and ferroptosis, we perturbed Drp-1 function by (1) shRNA mediated knock-down of Drp1 (Figure S4B-S4C) or by (2) over-expressing the dominant negative mutant Drp1-K38A (Drp1-K38A OE)^39^. In both approaches, Drp1 perturbation in confined cells inhibited mitochondrial fragmentation (Figure 4C-4F), while significantly reducing the upregulation of total and lipid ROS level resulting in a decrease in cell death (Figure 4G-4I). Taken together, these results indicate that in confined cells, Drp1 triggers mitochondrial fission which is associated with elevated mtROS levels in the confinement-induced ferroptosis cascade.

### 4. Nuclear confinement increases Drp1-phase separation

We next asked how Drp1 achieves spatio-temporal activation in cells under axial confinement. First, we examined whether ER constriction of mitochondria was triggering Drp1-dependent mitochondrial fission^40^. As the rough ER membrane is continuous with the outer nuclear membrane, we speculated that ER-mitochondria contacts could also be affected by nuclear deformation during axial confinement. To address this, we used the contact FP system^41^ which targeted dimerisation-dependent fluorescent proteins (ddFP) to organelle membranes to visualize organelle contact sites. Using this approach, we observed that the number of ER-mitochondria contacts remained unchanged even after mitochondrial fragmentation during confinement (Figure S5A-S5B). Thus we concluded that ER-mitochondrial contacts during confinement were not triggering Drp1 activation.

We next monitored endogenously tagged Drp1 spatio-temporal dynamics in confined cells via CRISPR-CAS9 knock-in of eGFP at the N-terminus of Drp1 locus (Figure S5C). Similar to a previous study^39^, endogenous Drp1 was mainly diffusive in the cytoplasm and occurred as puncta co-localized with mitochondria (Figure S5D). When cells were subjected to axial confinement we observed an increase in both the number of eGFP-Drp1 puncta as well as the association with mitochondria (82.75 % at 40 min vs. 74.42 % at 0 min) (Figure 5A-5C). Specifically, when we focused on mitochondrial fission sites, the majority exhibited eGFP-Drp1 puncta localization (90.79 %, Figure 5D, Movie S5). Curiously to us, these puncta were reminiscent of phase separated condensates. Experiments in line with this hypothesis demonstrated that when we subjected purified Drp1 to phase separation conditions, instantaneous droplet formation was seen to occur (Figure 5E-5F, Figure S5E-S5F).

**Figure 5.**
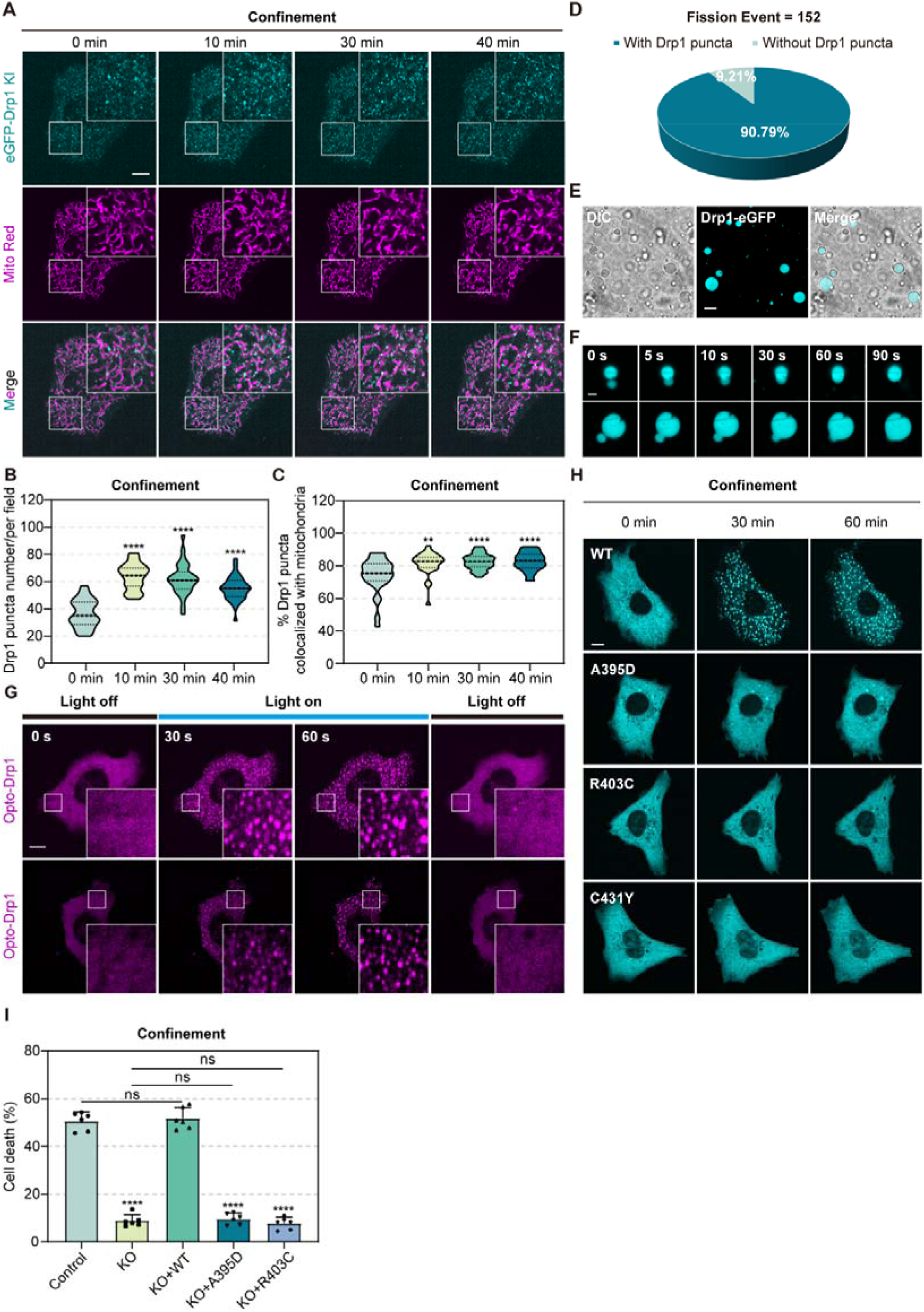
Nuclear confinement increases Drp1-phase separation. (A) Representative time-lapse images of endogenous eGFP-Drp1 knock-in HeLa cells co-stained with Mito Red during 40 min confinement. Insets are magnified images of areas indicated by white boxes. Scare bar = 10_μm. (B) Quantification of the number of Drp1 puncta per field of view in cells confined for 0 min, 10 min, 30 min, and 40 min (*N* = 30 fields from 18 cells, *\*\*\*\*p* < 0.0001). (C) Quantification of the proportion of Drp1 puncta localized to mitochondria in cells confined for 0 min, 10 min, 30 min, and 40 min (*N* = 30 fields from 18 cells, *\*\*p* < 0.01, *\*\*\*\*p* < 0.0001). (D) Pie chart showing the proportion of fission sites with a Drp1 puncta (*N* = 152 events from 7 cells). (E) Representative DIC and fluorescence images of purified Drp1-eGFP (20 μM) diluted in 200 mM NaCl containing 8% dextran. Scale bar = 10 μm. (F) Representative time-lapse images demonstrating small Drp1-eGFP droplets fusing to form larger droplets. Scale bar = 2 μm. (G) Representative images of HeLa cells expressing Opto-Drp1 in the absence and presence of blue light (off/on). Insets are magnified images of areas indicated by white boxes. Scale bar = 10 μm. (H) Representative time-lapse images of HeLa cells expressing eGFP-tagged wild-type Drp1 or Drp1 mutants (A395D, R403C, C431Y) during confinement. Scale bar = 10 μm. (I) The percentage of cell death in Control, Drp1 KO, KO+WT (Drp1 KO rescued with wild-type eGFP-Drp1), KO+A395D (Drp1 KO rescued with eGFP-Drp1 A395D mutant) and KO+R403C (Drp1 KO rescued with eGFP-Drp1 R403C mutant) HeLa cells upon confinement (*N* = 6 fields, ns represents not significant, *\*\*\*\*p* < 0.0001).

To determine whether similar phenomena occur in cells undergoing axial confinement, we employed an optogenetic approach based on the optoDroplet system^42^. Our opto-Drp1 system allows for the reversible formation of droplets via tagged CRY2 to Drp1 upon blue light exposure (Figure 5G), suggesting that the Drp1-puncta and droplets we observe are potentially mediated by liquid-liquid phase separation (LLPS). To extensively understand which domains of Drp1 are responsible for its phase separation ability, we used a series of disease related Drp1 point mutants (A395D, R403C, C431Y)^43^ shown to abrogate Drp1 function, hypothesizing that the abrogated Drp1 function of these mutants could be a result of their ability to phase separate. Consistent with our hypothesis, the expression of various point mutants in the middle domain of Drp1 abrogated the formation of Drp1 droplets in cells and in-vitro system (Figure 5H, Figure S5G). Supporting this, we observed that the formation of these puncta within cells was not dependent on the GTPase domain (Figure S5H). Importantly, these mutants, when expressed in Drp1 knock-out cells (Figure S5I) were capable of resisting cell death induced by axial confinement (Figure 5I). Together, these data show that axial confinement induces endogenous Drp1 puncta formation, localizing to mitochondria as a likely consequence of Drp1 phase separation.

### 5. mtROS increase and cPLA2 activation concertedly trigger ferroptosis under nuclear confinement and in osteoarthritis

While our observation of mitochondrial fragmentation occurred prior to the mitochondrial translocation of cPLA2 (Figure 2H, Movie S3), we wondered if, and how, these two events were mechanistically linked. To understand this, we monitored cPLA2 localization in Drp1-depleted cells and found that cPLA2 could still translocate to mitochondria under confinement, even when mitochondrial fragmentation was abolished by Drp1-depletion (Figure 6A). This finding suggests that mitochondrial translocation of cPLA2 is not due to mitochondrial fragmentation. Supportive of this finding, cPLA2 inhibition in confined cells did not prevent mitochondrial fragmentation (Figure S6A-S6B) nor the total ROS generation from increasing (Figure S6C-S6D). Finally, treating cells with increasing concentrations of arachidonic acid (ARA), which is released downstream of cPLA2, had negligible impacts on lipid ROS level and cell viability (Figure S6E-S6F), suggesting that ferroptosis cannot be induced solely by effects of cPLA2 translocation and activation.

**Figure 6.**
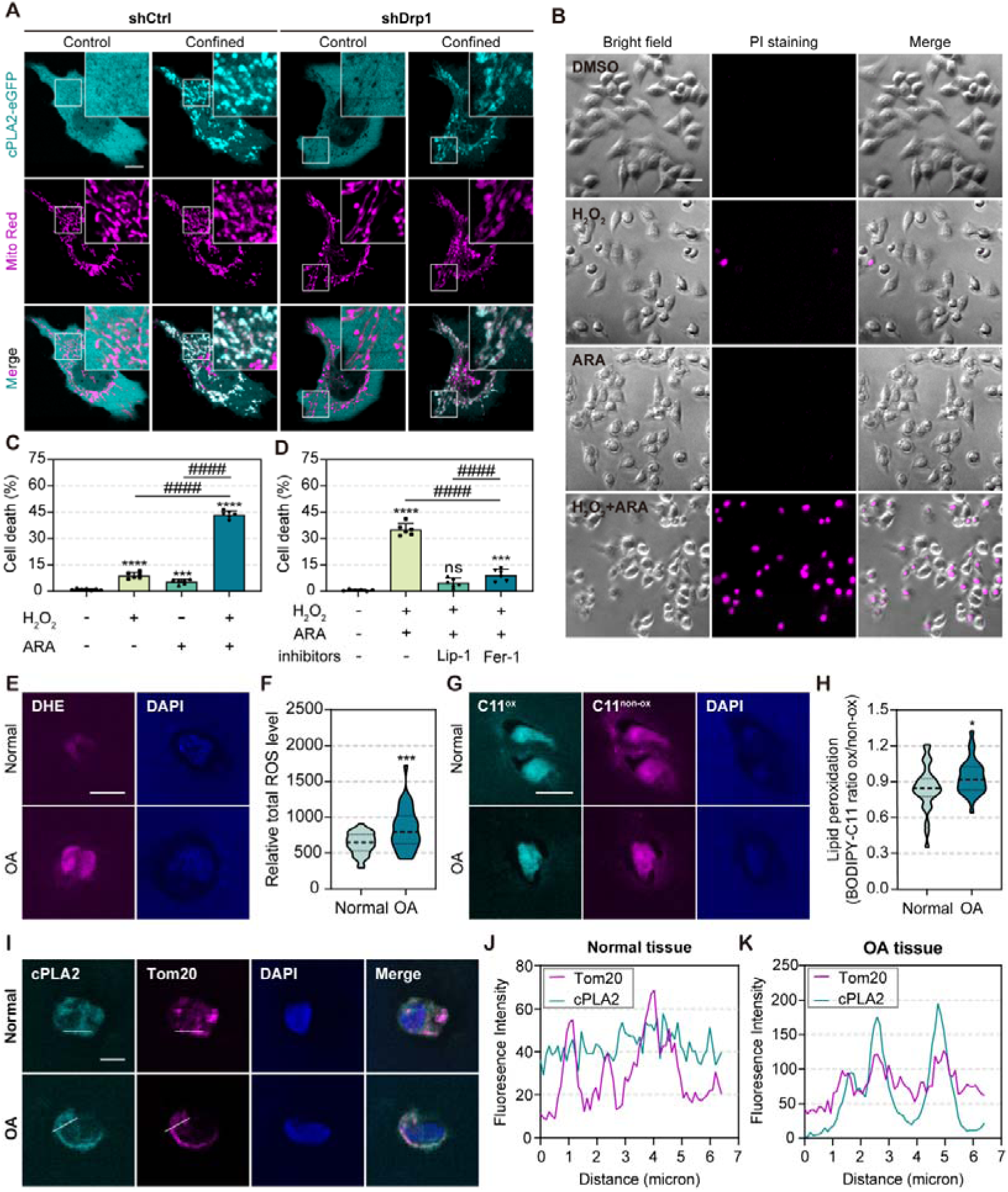
mtROS increase and cPLA2 activation concertedly trigger ferroptosis under nuclear confinement and in osteoarthritis. (A) Representative live-cell images of the cPLA2-eGFP expressing shCtrl or shDrp1 HeLa cells co-stained with Mito Red upon confined and unconfined conditions. Scale bar = 10 μm. (B) Representative bright field and fluorescence images of PI staining in HeLa cells treated with DMSO, H_2_O_2_ (400 μM) alone, ARA (400 μM) alone, and H_2_O_2_ + ARA. Scale bar = 40 μm. (C) Quantification of the percentage of cell death in HeLa cells related to B (*N* = 6 fields, *\*\*\*\*p* < 0.0001, ^####^*p* < 0.0001). (D) The percentage of cell death in HeLa cells treated with DMSO, H_2_O_2_ (400 μM) + ARA (400 μM), H_2_O_2_ + ARA + Lip-1 (10 μM), and H_2_O_2_ + ARA + Fer-1 (10 μM) (*N* = 6 fields, ns represents not significant, *\*\*\*p* < 0.001, *\*\*\*\*p* < 0.0001, ^####^*p* < 0.0001). (E) Representative images of DHE staining (50 μM for 2 hours) in relative normal and OA tissues. The relative normal tissue and OA tissue are from the same patient. Total ROS (DHE, magenta), Nuclei (DAPI, blue). Scale bar = 10 μm. (F) Quantification of total ROS level in normal and OA tissue related to E (*N*_Normal_ = 36 cells, *N*_OA_ = 34 cells, *\*\*\*p* < 0.001). (G) Representative images of BODIPY-C11 581/591 staining (40 μM for 2 hours) in normal and OA tissues. Non-oxidative state (magenta), Oxidative state (cyan), Nuclei (DAPI, blue). Scale bar = 10 μm. (H) Quantification of lipid ROS level in normal and OA tissue related to G (*N* = 30 cells, *\*p* < 0.05). (I) Representative images of cPLA2 and Tom20 in normal and OA tissue. cPLA2 (cyan), Tom20 (magenta), Nuclei (DAPI, blue). Scale bar = 5 μm. (J-K). Line scan plot showing the distribution of cPLA2 and Tom20 along the direction of the white arrowheads related to I in normal (J) and OA tissue (K).

While it is understood that the generation of lipid hydroperoxides from ARA requires cellular ROS^31^, we wondered whether it was possible to mechanistically recapitulate the confinement-induced ferroptotic cell death from the generation of lipid hydroperoxides by exposing cells to ARA in tandem with ROS, like H_2_O_2_. Indeed, only when cells were exposed to this combination of treatments, and not individually, we observed a potent induction of cell death (Figure 6B-6C), which were antagonized by treating cells with the ferroptosis inhibitors including Liproxstatin-1 and Ferrostatin-1 (Figure 6D).

To understand whether our mechanistic findings could be applied to an outstanding clinical question, we turned to the pathphysiological process of osteoarthritis (OA). In OA excessive mechanical loading of chondrocytes, inflammation, cell death and ferroptosis are reported to occur^18^. Thus, we asked whether ferroptosis in OA was triggered by the same molecular cascade of events that we observed in-vitro. Using normal and OA human cartilage samples, we compared total ROS and lipid ROS levels and observed higher levels of total (Figure 6E-6F) and lipid ROS (Figure 6G-6H) in OA regions. Further, immunostaining of cPLA2 and Tom20 in chondrocytes of OA tissue revealed partial co-localization (Figure 6I-6K). Our findings suggest that the same molecular players we find responsible for confinement-induced ferroptosis in-vitro are also present clinically in OA.

## Discussion

Here, we show that prolonged axial confinement of non-motile adherent cells results in cell death via ferroptosis. In clinically relevant patient samples, these findings provide new insight to the clinically relevant pathological processes of OA. We show that when cells are mechanically confined, they can undergo ferroptosis driven principally by mitochondrial fragmentation and cPLA2 activation. This mechanically activated pathway, disrupts mitochondrial homeostasis. While our observation of mitochondria fragmentation during confinement is partly explained by Drp1-dependent fission, perplexingly we did not observe evidence of increased ER-Mitochondria contact sites that incline to recruit Drp1^44^. One possible explanation could be that while ER-mitochondrial contacts remain stable in number, other organelle-mitochondrial membrane contact sites could increase, recruiting mitochondrial division machinery. An example of this could occur through the interaction between mitochondria and lysosomes, which has been shown to induce mitochondrial fission^45^. The other possibility is that increased fission events during confinement may still be evoked by the pre-existing ER-mitochondria contacts, possibly triggered by calcium transfer between these two organelles^46^, known to regulate mitochondrial fission.

Our understanding of Drp1 as the master executor of mitochondrial membrane scission through the recruitment by the adaptor proteins followed by self-assembling into helix^38, 39^, is well characterized. However, how these protein are regulated and coordinated biochemically and biophysically to assemble division machinery to drive mitochondrial fission is lesser known. Our study reveals that Drp1 is capable of undergoing phase separation in cells upon confinement, which we demonstrate via endogenous Drp1 tagging. Curiously, when utilizing the optoDroplet system to induce Drp1 foci formation, we found that when we co-expressed the eGFP tagged cytoplasmic adaptor proteins MFF or MID49/51, these adaptor proteins also condensed into droplets that co-localized with Drp1 puncta (Data not showns here). These results suggest that the Drp1 may assemble into complexes with adaptor proteins required for mitochondrial fission via phase separation.

Our work shows that the nucleus act as the central mechanosensor in confinement-induced mitochondrial fission, resulting in ferroptosis. Whilst we show insight to the molecular transduction of nuclear deformation, leading to mitochondrial fragmentation, it will be important to investigate the possible role of the that organelle-mitochondrial membranes contact sites may play. While some membrane contacts are better studied (such as the ER contact sites), others like mitochondria-nuclear membrane contacts have not been well described in detail, which could explain our findings. Although changes in nuclear shapes often directly modulate gene expressions in cells^47^, and over 95% of mitochondrial proteins are encoded by nuclear DNA^48^, it is unlikely that the nucleus regulates mitochondrial fragmentation in confinement at the transcriptional or translational levels as fragmentation occurs early during confinement, (30 to 60 min). Thus, further research to elucidate the underlying interaction between the nucleus and mitochondria will shed light on the ruler roles of nucleus in regulating cell biological processes under confinement.

Our data identify a new role for nuclear deformation in mechano-induced ferroptosis-dependent cell death, which leads to inflammation. Aberrant mechanical stimuli usually lead to traumatic events in cells and are linked to several diseases like cancer and OA^18, 49^ which are inflammatory conditions. Our data implicate that nuclear deformation-mediated mitochondrial dysfunction coordinates cPLA2 to trigger ferroptosis under mechanical confinement, which may be involved in OA progression. Our findings may provide a framework for further comprehensive understanding and targeting of ferroptosis in non-only treating OA but may also give insight to other diseases related to mechanical stresses and inflammation.

## Method

### 1.1 Cell culture

Human cervical carcinoma HeLa cells were a generous gift from Dr. Yong Liu (Xuzhou Medical University). Human embryonic kidney 293T (HEK293T) cells were a generous gift from Dr. Yuxin Yin (Peking University). Cells were cultured in Dulbecco’s modified Eagle medium (DMEM; Corning, 10-013-CRVC) supplemented with 10% fetal bovine serum (FBS; PAN-Biotech, P30-3302), 100 U/mL penicillin and 100 μg/mL streptomycin (Macgene, CC004) at 37°C with 5% CO_2_. For passage, the cells were washed with PBS (Macgene, CC010) and detached by trypsin-EDTA (Macgene, CC012). For long-term live cell imaging, cells were plated on fibronectin (Sigma-Aldrich, F1056-5MG, 10 μg/mL)-coated 35 mm glass-bottom dishes (Cellvis, D35-10-1-N) and maintained in CO_2_-independent DMEM (Gibco, 18045-088) supplemented with 10% FBS, 100 U/mL penicillin and 100 mg/mL streptomycin at 37°C throughout the imaging process.

### 1.2 Plasmid construction and DNA transfection

The full-length human Drp1 was cloned using plasmid containing Drp1 from Dr. Yuhui Zhang (Huazhong University of Science and Technology) and subcloned into a lentiviral vector (pLVX-AcGFP-N1) with N-terminal tagged eGFP using an ABclonal MultiF Seamless Assembly kit (ABclonal, RK21020). The Drp1 truncations and single-point mutations were amplified and cloned into the same vectors. The FL-Drp1 (1-699 aa), GTPase domain (1-302 aa), ΔGTPase domain (303-699 aa), FL-Drp1-A395D, FL-Drp1-R403C and FL-Drp1-C431Y were constructed. The FL-Drp1-K38A was cloned from the pcDNA-Drp1K38A plasmid, a gift from Alexander van der Bliek & Richard Youle (Addgene plasmid # 45161 ; http://n2t.net/addgene:45161 ; RRID:Addgene_45161), and subcloned into pLVX-AcGFP-N1 with N-terminal tagged mCherry. Plasmid containing Cry2 was a gift from Chandra Tucker (Addgene plasmid # 60032 ; http://n2t.net/addgene:60032 ; RRID:Addgene_60032) to generate the Opto-droplet Optogenetic system (Opto-Drp1). The human cPLA2 was cloned from the HeLa cell-extracted cDNA library and subcloned into pLVX-AcGFP-N1 with C-terminal tagged eGFP. The Contact-FP constructs (GA and GB) were cloned from GA-NES (a gift from Robert Campbell; Addgene plasmid # 61018 ; http://n2t.net/addgene:61018 ; RRID:Addgene_61018) and GB-NES (a gift from Robert Campbell; Addgene plasmid # 61017 ; http://n2t.net/addgene:61017 ; RRID:Addgene_61017), and were subcloned into the pLVX-AcGFP-N1 with C-terminal tagged full-length sequence of Sec61β (pLVX-GA-Sec61β) or mitochondrial localization sequence of MAVS (pLVX-GB-MAVS), respectively.

For transient transfection, HeLa cells seeded on 6-well plates were transfected with 2 μg midi-prep quality plasmid DNA in Opti-MEM (Invitrogen, 31985-070) containing 2 μL Neofect™ DNA transfection reagent (TF20121201) following the protocol, for 24-48 h. For viral particle generation, HEK293T cells seeded on 10 cm culture dishes (Nest, 704201) were co-transfected with 5 μg of the expression plasmid, with 3.75 μg packaging plasmids psPAX2, and 1.25 μg pCMV-VSV-G, in Opti-MEM containing 10 μL Neofect™ DNA transfection reagent.

### 1.3 Antibody and Reagents

The following antibodies were used in this study: rabbit anti-Drp1 (EPR19274, dilution 1:1000 for Western blotting), rabbit anti-cleaved N-terminal GSDMD (EPR20829-408, dilution 1:1000 for Western blotting), rabbit anti-GSDME (EPR19859, dilution 1:1000 for Western blotting), rabbit anti-MLKL (EPR9514, dilution 1:1000 for Western blotting), rabbit anti-GAPDH (EPR16891, dilution 1:4000 for Western blotting) and mouse anti-β-actin (ab8226, dilution 1:2000 for Western blotting) were purchased from Abcam; Rabbit anti-Drp1 (A21968, dilution 1:1000 for Western blotting), rabbit anti-full-length GSDMD (A17308, dilution 1:1000 for Western blotting), rabbit anti-cPLA2 (A0394, dilution 1:1000 for Western blotting), rabbit anti-Tom20 (A19403, dilution 1:200 for immunofluorescence staining) and mouse anti-α-tubulin (AC012, dilution 1:3000 for Western blotting) were purchased from Abclonal; Mouse anti-cPLA2(sc-454, dilution 1:100 for immunofluorescence staining), anti-mouse (sc-516102, dilution 1:4000) and anti-rabbit (sc-2004, dilution 1:2000) horseradish peroxidase (HRP)-conjugated secondary antibodies were from Santa Cruz Biotechnology. Anti-mouse Alexa Fluor 488 (A-21202) and anti-rabbit Alexa Fluor 555-conjugated secondary antibody (A-31572) were from ThermoFisher Scientific.

For commercial reagents, Z-VAD-FMK (S7023), DFOM (S5742), Liproxstatin-1 (S7699), Ferrostatin-1 (S7243), and MitoTEMPO (S9733) were from Selleckchem. Trolox (ab120747) was from Abcam. FCCP (HY-100410), CAY10650 (HY-10801), and Arachidonic acid (HY-109590) were from MedChemExpress. NSA was a gift from Dr. Huang Wang (Peking University). TNFα was a gift from Dr. Fuping You (Peking University). Smac was a gift from Dr. Xiaodong Wang (National Institute of Biological Sciences, Beijing). MitoTracker^TM^ Red CMXRos (M7512), Mito Tracker^TM^ Green FM (M7514), TMRM (T668), and Prolong^TM^ Diamond Antifade with DAPI (P36962) were from ThermoFisher Scientific. Other drugs used in this study included EUK134 (TargetMol, T6495), NAC (Shanghai Yuanye Bio-Technology, S20137), and H_2_O_2_ (Aladdin, H112515).

### 1.4 Cell confinement

Two cell confinement system allowed for the monitoring of multiple fields of live cells over the entire duration of prolonged axial confinement were used in our study. The first one was a bespoke, in-house device consisting of stainless-steel weights, optical glass, and microsphere spacers of different sizes (Bangs Laboratories, USA) (Figure S1B). For biochemical studies, the mixture of cells and microspheres with appropriate volume was plated into the 35 mm culture dish (Corning, 430165) and incubated overnight. Then, weighted optical glass was carefully placed on top of the culture substrate. The diameters of the microspheres (20 μm versus 3 μm) determined the height for spatial confinement of cells between the glass and the substrate. Notably, a version of the cell confiner adapted to 35 mm glass-bottom dishes was used to perform high-resolution live cell imaging.

The second one were composed of a custom designed PDMS (Slygard 184, Dow Corning) piston, as previously described^19^. The degree for spatial confinement of cells was determined by a coverslip containing 3 µm height PDMS micropillars, which were fabricated following standard procedures and were passivated with non-adhesive pLL(20)-g[3.5]-PEG(2) (SuSoS).

### 1.5 Microscopy imaging of cell death

Cell death in confinement was evaluated by propidium iodide (PI, Sigma-Aldrich, P4170-10MG) uptake of cells measured by fluorescence imaging. PI was then added to the culture medium at a final concentration of 5 μg/mL before confinement. Static images of PI positive cells were captured using a Plan Apo 4x air objective lens on a Spinning Disk Confocal Microscope (Andor Dragonfly) after 8-10 hours of confinement. Time-lapse imaging of cell death was imaged via epifluorescence microscopy (LIFE EVOS FL Auto fluorescent microscope, Life Technologies, USA). Raw full bit depth images were processed using ImageJ software (https://imagej.nih.gov/ij/). Image data shown were representative of at least three randomly selected fields.

### 1.6 Measurement of ROS

To detect total ROS, cells were pre-stained with 10 μM DHE (Beyotime, S0063) in culture media at 37°C for 30 min, before washing twice with PBS. To detect lipid ROS, cells were pre-incubated with 5 μM BODIPY-C11 581/591 staining (Abclonal, RM02821) at 37°C for 1 h and washed twice with PBS. After labelling, cells were subjected to confinement and fluorescence images were taken by a 10× air objective lens of CellTIRF microscopy (Olympus, Japan) at different time points. To detect mtROS, cells were stably expressed with Mito-roGFP, and imaging was acquired using a 63× objective lens on Andor Dragonfly Spinning Disk Confocal Microscope with the excitation wavelength of 405 nm or 488 nm, and the emission wavelength of 500-550 nm. mtROS level was indicated by the RoGFP ratio, which was calculated as the fluorescence intensity under the excitation wavelength of 405 nm divided by the intensity under the excitation wavelength of 488 nm (405/488 Ratio).

### 1.7 Western blot

For Western blot, cells were washed with PBS and lysed with appropriate volumes of commercial 1× SDS-PAGE sample loading buffer (Beyotime, P0015A) for 5-10 min. The protein samples were boiled for 15 min at 95°C and were separated on SDS-PAGE gels before being transferred onto nitrocellulose membranes by wet electrophoretic transfer. After blocking in PBS containing 5% milk, blots were incubated with primary antibodies at 4°C overnight or for 2 h at room temperature, followed by the secondary antibody incubation for 1 h at room temperature. Blots were visualized and recorded using SH-Compact 523 (SHST). The images were processed by ImageJ software.

### 1.8 Quantitative real-time PCR

Briefly, total RNA from HeLa cells were isolated using Trizol (Life Technologies, 15596026). RNA was reverse transcribed using a Transcript One-Step gDNA Removal and cDNA Synthesis SuperMix Kit (Transgene, AT311-02). Level of the ASCL4, COX2 and α-tubulin genes were analyzed by qRT-PCR amplified using SYBR Green Fast qPCR Mix (ABclonal, RK21203). Data shown were the relative abundance of the ASCL4 and COX2 mRNA normalized to α-tubulin.

The primer sequences are listed below.

ACSL4 - F: 5′-GCTATCTCCTCAGACACACCGA-3′;
ACSL4 - R: 5′-AGGTGCTCCAACTCTGCCAGTA-3′;
COX2 - F: 5′-CGGTGAAACTCTGGCTAGACAG-3′;
COX2 - R: 5′-GCAAACCGTAGATGCTCAGGGA-3′;
α-tubulin - F: 5′-CGGGCAGTGTTTGTAGACTTGG-3′;
α-tubulin - R: 5′-CTCCTTGCCAATGGTGTAGTGC-3′.

### 1.9 Cellular enucleation

Enucleation was performed as previously described^34^ with some modifications. Briefly, small plastic slides that fit into 1.5 mL Eppendorf tubes were cut from 10 cm cell culture dishes, followed by sterilisation in 95 % ethanol for 10 min followed by ultraviolet irradiation for 30 min. Before seeding cells, the plastic slides were coated with a 10 μg/mL fibronectin (Sigma-Aldrich, F1056-5MG, 10 μg/mL) solution for 30 min at 37°C. Cells on plastic slides were pre-incubated with DMEM containing 2 μM Cytochalasin D (CytD, Sigma-Aldrich, C2618-200UL) in 1.5 mL Eppendorf tubes for 1 h at 37°C, then the cells were centrifuged at 16,000 g in a GeneSpeed microcentrifuge (Gene Company Limited, 1524) for 30 min. After centrifugation, the remaining cells and cytoplasts on the plastic slides were placed in 10 cm cell culture dishes containing fresh medium without CytD to spread and recover from the effects of CytD fully. About 3 to 4 hours later, the cells were washed with PBS, trypsinised, and reseeded on fibronectin-coated glass-bottom dishes for live cell imaging.

### 1.10 Mitochondrial Morphodynamics Analysis

Segmentation of the mitochondrial outer membrane (Tom20-mCherry signal) was done using the auto-context workflow from Ilastik^50^. First, raw images were converted into HDF5 files and imported in Ilastik. For the first round of training, the features selected were color/intensity, edge and texture. The σ value for all features was set to 0.7px. The algorithm was trained for two labels: mitochondrial outer membrane and background. As minor bleaching occurred during the acquisition, randomly chosen images were selected throughout the entire acquisition to account for visually different parts of the dataset. For the second round of the auto context workflow, the σ values for all features were set to 0.7px. Then, segmentation was performed identically as during the first round. After simple segmentation predictions were deemed satisfactory, the simple segmentation masks were exported as multipage tiff files. These multipage tiff files were then imported to ImageJ and despeckled using the despeckle function. Images were then inverted and binarized before particles analysis. Datasets were binned into the 10 first frames (first 20 minutes) and the following 35 frames (last 70 minutes of acquisition). Size threshold was 9-150 (Pixel units) and circularity 0.3-1.0 to exclude areas between distinct mitochondria for the first 10 frames. Size and circularity thresholds were set to 9-600 (Pixel units) and 0.3-1.0 for the second part of the acquisition. Regions of interests (ROIs) around individual cells were drawn for each timepoint and particles for each cell were analysed. Average perimeter, average circularity, and total number of particles per cell were calculated and plotted in a XY graph. Data was imported and plotted using Graphpad Prism version 9.5.1 for Windows, GraphPad Software, Bosoton, Masschusetts USA, www.graphpad.com.

### 1.11 Atomic force microscopy (AFM) coupled with live cell imaging

To confine a single cell, we used a Bioscope Resolve AFM (Bruker, USA) with the HA_FM tipless silicon probe (force constant, 3 or 6 N/m, ScanSens), and the AFM was mounted on the stage of an inverted fluorescence microscope (ECLIPSE Ti, Nikon) for imaging. Before confinement and imaging, HeLa cells were plated on fibronectin-coated 35 mm glass-bottomed dishes (Cellvis, D35-20-1-N). Experiments were initiated 4-6 h after cell plating to allow for cells to fully spread. During imaging, cells were maintained in CO2-independent DMEM supplemented with 10% FBS, 100 U/mL penicillin, and 100 mg/mL streptomycin. The probe was mounted in the AFM head and the 6 N/m microcantilever was lowered on the cell until a 0.8 V setpoint was reached, at which cells were significantly confined. Meanwhile, fluorescence images were taken every 0.2 seconds using a 100× oil immersion objective to record the whole process of microcantilever lowering. After cells were stably compressed by the cantilever, images were taken every 30 seconds to observe changes in mitochondrial morphology. All microscopy equipment was placed in an acoustic enclosure on the top of the vibration isolation platform.

### 1.12 Generation of eGFP-Drp1 Knock-in cells

HeLa cell line with an endogenous N-terminal eGFP insertion in the Drp1 gene locus was generated via the CRISPR/Cas9 system. The following sgRNA was generated via the online CRISPR design tool (CRISPR tefor.net) and cloned into a lentiCRISPR-V2 vector (a gift from Feng Zhang; Addgene plasmid # 52961 ; http://n2t.net/addgene:52961 ; RRID:Addgene_52961). The donor template contained a left homologous arm (840 bp before the start code), eGFP sequence, and right homologous arm (1000 bp after the start code). The homologous arms were amplified from the HeLa cell-extracted genome DNA. Cells were co-transfected with sgRNA and relative donor template plasmid for 24 h and then treated with puromycin (Sigma-Aldrich, P8833) at 2 μg/mL for 24 h. Surviving cells were sorted by Fluorescence Activated Cell Sorting (FACS) to isolate fluorescence-positive cells. The candidate single cell clones were cultured and verified by Western blot.

sgRNA for Drp1 N-terminal knock-in in HeLa cells:

sgRNA: 5’-CGCCGGCCACGGCAATGAAT-3’.

### 1.13 Protein expression and purification

For recombinant protein expression in *E.coli* BL21-CondonPlus (DE3)-RIPL (AngYu Biotechnologies, AYBIO-G6028), DNA fragments encoding Drp1 were cloned into the modified pET28a (+) vector with an N-terminal His6-tag and a C-terminal eGFP tag as described previously (Liang et al., 2020). Bacteria transformed with plasmid were cultured in LB medium with 50 mg/L kanamycin at 37°C to an OD600 of 0.8-1.0. After that, protein expression was induced with 0.1 mM isopropyl-1-thio-β-D-galactopyranoside (IPTG) for 16 h at 18°C. For protein purification, cell pellets were collected by centrifugation at 4000 g and were suspended in TBS buffer containing 20 mM Tris (pH 8.0), 140 mM NaCl, 30 mM KCl, 10% glycerol, 20 mM imidazole, 0.5 mM TCEP, and protease inhibitor mixture (Complete EDTA-free, Roche). The suspension was lysed by sonication and cleared by centrifugation twice at 4°C,13000 g for 30 min. The supernatant was applied to a Ni-IDA column (Smart-Lifesciences) and washed three times using buffers containing 20 mM Tris (pH 8.0), 500 mM NaCl, 0.5 mM TCEP, and different concentrations of imidazole (0 mM, 20 mM, 40 mM). Bound Drp1 proteins were eluted with buffer containing 20 mM Tris (pH8.0), 500 mM NaCl, 0.5 mM TCEP and 300 mM imidazole, and PreScission protease was added to the elution to cleave the N-terminal His_6_-tag. Finally, proteins were purified by size exclusion chromatography using Superose 6 Increase 10/300 (GE Healthcare) in buffer containing 20 mM Tris (pH 8.0), 500 mM NaCl and 0.5 mM TCEP. Fractions containing Drp1 were collected, concentrated, and flash frozen in liquid nitrogen. Protein concentrations were measured with NanoDrop 2000 spectrophotometry using the theoretical molar extinction coefficients at 280 nm. Protein purity was evaluated with Coomassie brilliant blue staining of SDS-PAGE gels.

Quality control data of the recombinant Drp-eGFP proteins were shown in Figure S5E-S5F.

### 1.14 In vitro phase transition assay

#### Preparing coverslips and glass slides

Glass slides were rinsed with ethanol twice and dried with air. A flow chamber for imaging with two openings on opposite sides was then made by attaching a coverslip (Fisherbrand, 12541016) to the glass slide using double-sided tape placed parallel to each other.

#### Phase separation assays

Full-length Drp1 and its mutants were labeled by eGFP tags. The purified proteins were saved in a high salt solution containing 500 mM NaCl, 20 mM Tris (pH = 8.0), and 0.5 mM TCEP after protein production. According to different protein and NaCl concentrations for phase separation assay, they were diluted into the corresponding lower salt experimental buffers at a final concentration of 8% dextran to induce phase separation. The protein-phase buffer mixtures were mixed gently. Then, 5 μL mixtures were injected into the homemade flow chamber for fluorescent imaging using the Andor Dragonfly confocal imaging system by putting the chamber upside down on the objective. Notably, the last layer (closest to the lens) of droplets was averted when acquiring the images to minimize the interaction between droplets and the glass surface.

### 1.15 Generation of knock-down and knock-out cells

For Drp1 and cPLA2 knockdown in HeLa cells, the related shRNA were cloned into pLKO.1 vector. After lentivirus infection and puromycin selection, the pooled cells were collected and verified by Western blot.

For Drp1 knockout, sgRNA was cloned into the LentiCRISPR-V2 vector, followed by transfection into HeLa cells. After puromycin selection, the single cell clones were cultured and verified by RNA sequence and Western blot.

The primes used to knockdown are listed below:

shscramble: 5’-AACGCTGCTTCTTCTTATTTA-3’;
shDrp1: 5’-GCTACTTTACTCCAACTTATT-3’;
ShcPLA2: 5’-CCTTGTATTCTCACCCTGATT-3’.
sgRNA for Drp1 knockout in HeLa cells:
sgRNA: 5’-AAATAGCTACGGTGAACCCG-3’.

Sequencing primer:

Drp1-seq-F: 5’- GATGCATCACATCAGAGATTGTT-3’;
Drp1-seq-R: 5’- TTGCCAAAGTGAAGGTGTCG-3’.

For integrin knockdown, the siRNAs targeting to different subunits (AV/A1/A3/A5/A8/B1/B3/B5/B6/B8) were purchased from Sangon Biotech.

### 1.16 Immunofluorescence staining of human cartilage samples

Human cartilage samples were obtained from Peking University Third Hospital. The relative normal and OA cartilage samples from the same patient with OA were frozen in OCT and sectioned into slices attached onto glass slides. Slides were washed by PBS for 5 min and were incubated with PBS containing 10% FBS and 0.3% Triton X-100 for 1 h at room temperature. Then, the slides were incubated with the primary antibodies mix solution containing mouse anti-cPLA2 (1:100), rabbit anti-Tom20 (1:200) and 0.3% Triton X-100 for 2 h at room temperature. After three times of PBS washing for 5 min each, the slides were incubated with anti-mouse Alexa Fluor 488- and anti-rabbit Alexa Fluor 555-conjugated secondary antibodies diluted in 1:200 for 2 h at room temperature. The secondary antibodies mix solution also contained 0.3% Triton X-100. Following another three times washing with PBS, a coverslip was mounted using with ProLong™ Diamond with DAPI. Images were captured on an Andor Dragonfly confocal imaging system after mounting medium was solidified.

To detect total ROS and lipid ROS levels in normal and OA cartilage samples, the slides were washed by PBS for 5 min and stained with DHE probe (50 μM) or BODIPY-C11 581/591 probe (40 μM) for 2 h at 37°C. Following staining, slides were washed three times with PBS for 5 min each and coverslips were mounted using with ProLong™ Diamond with DAPI.

Above procedure was conducted according to guidelines approved by Medical Science Research Ethics Committee of Peking University Third Hospital (ethical approval No.: 2013003). Informed consent was obtained from subject and the experiments were conformed to the principles set out in the WMA Declaration of Helsinki and the Department of Health and Human Services Belmont Report.

### 1.17 Statistics and Reproducibility

All data were representative of two or more independent experiments and were expressed as mean ± SD (standard deviation). Unless stated otherwise, statistical analysis was determined using two-tailed unpaired or paired Student’s t-test for two groups comparison, or one-way ANOVA analysis with Tukey’s test for more than two group comparison, or two-way ANOVA analysis with Sidak’s test for multiple pairwise comparisons. All calculations of significance were performed with GraphPad Prism 8.0 Software.

## Extended Figures

**Figure S1.**
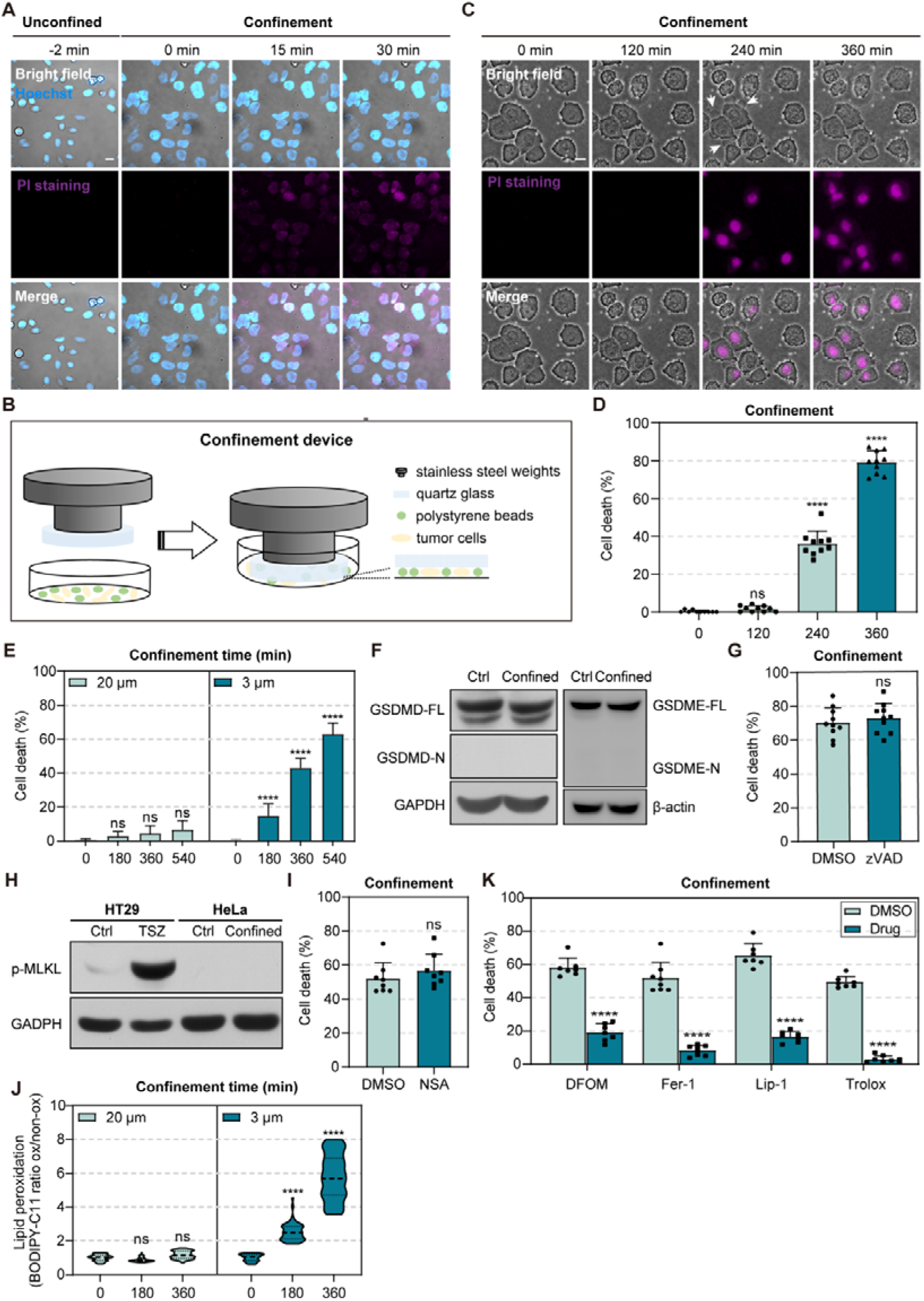
Confinement-induced cell death is not apoptosis, pyroptosis or necroptosis. Related to Figure 1. (A) Representative time-lapse images of HeLa cells beneath the PDMS micropillars during confinement in the presence of PI (magenta, 10 μM). Nuclei labeled with Hoechst (blue). Scale bar = 20 μm. (B) Schematic diagram of the second axial confinement device. By combining stainless steel weights, quartz glass, and polystyrene microspheres of different sizes, we can vertically restrict cell height through sandwiching cells between glass and culture dish, and polystyrene microspheres of defined diameters act as spacers to control the confinement height. (C) Representative time-lapse images of HeLa cells during confinement in the presence of PI. White arrow: large plasma membrane blebs. Scale bar = 20 μm. (D) The percentage of cell death in HeLa cells during confinement (*N* = 10 fields, ns represents not significant, *\*\*\*\*p* < 0.0001). (E) The percentage of cell death in HeLa cells during 20 μm or 3 μm confinement (*N* = 6 fields, ns represents not significant, *\*\*\*\*p* < 0.0001). (F) Western blot does not show the cleavage of GSDMD or GSDME in HeLa cells upon confinement (FL, full-length, N, N-terminal). GAPDH or β-tubulin is used as the loading control. (G) The percentage of cell death in HeLa cells treated with DMSO or pan-caspase inhibitor Z-VAD-FMK (100 μM) (*N* = 10 fields, ns represents not significant). (H) Western blot shows non-detectable phosphorylation of MLKL in HeLa cells upon confinement. HT29 cells treated with TNFα (T, 20 ng/mL), Smac (S, 100 μM), and Z-VAD-FMK (Z, 20 μM) for 8 hours act as positive control of necroptosis. GAPDH is used as the loading control. (I) The percentage of cell death in HeLa cells treated with DMSO or necroptosis inhibitor NSA (10 μM) upon confinement (*N* = 8 fields, ns represents not significant). (J) Lipid ROS level in HeLa cells upon 20 μm or 3 μm confinement (*N* = 50 cells, ns represents not significant, *\*\*\*\*p* < 0.0001). (K) The percentage of cell death in HeLa cells treated with DMSO or ferroptosis inhibitors upon 3 μm confinement mediated by microsphere spacers, including DFOM (100 μM), Ferrostatin-1 (Fer-1, 1 μM), Liproxstatin-1 (Lip-1, 0.5 μM), and Trolox (200 μM) (*N* = 7 or 8 fields, *\*\*\*\*p* < 0.0001).

**Figure S2.**
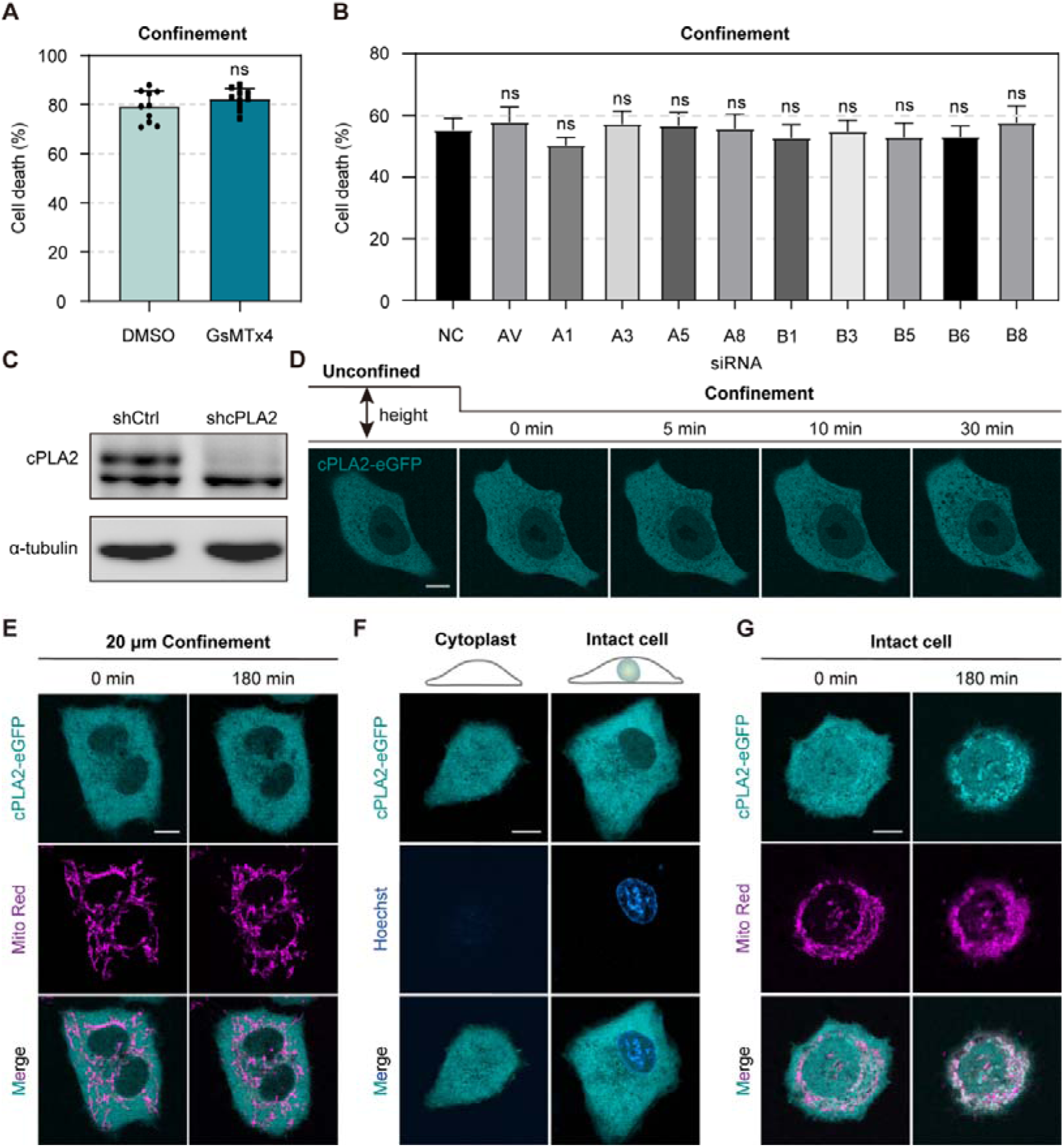
Nuclear confinement is critical for cPLA2 translocation to mitochondria. Related to Figure 2. (A) The percentage of cell death in HeLa cells treated with DMSO or GsMTx4 (10 μM) upon 3 μm confinement (*N* = 10 fields, ns represents not significant). (B) The percentage of cell death in HeLa cells transfected with siRNA targeting different integrin subunits upon 3 μm confinement (*N* = 8 fields, ns represents not significant). (C) Western blot shows the protein level of cPLA2 in shCtrl and shDrp1 HeLa cells. α-tubulin is used as the loading control. (D) Representative time-lapse images of cPLA2-eGFP expressing HeLa cells during 3 μm confinement. Scale bar = 10 μm. (E) Representative live-cell images of the cPLA2-eGFP expressing HeLa cells co-stained with Mito Red upon 20 μm confinement. Scale bar = 10 μm. (F) Representative live-cell images of cPLA2-eGFP expressing intact HeLa cells and enucleated cytoplasts. Nuclei labeled with Hoechst (blue). Scale bar = 10 μm. (G) Representative live-cell images of cPLA2-eGFP intact cells co-stained with Mito Red upon 3 μm confinement. These intact cells undergo the same treatment as enucleated cytoplasts before confinement. Scale bar = 10 μm.

**Figure S3.**
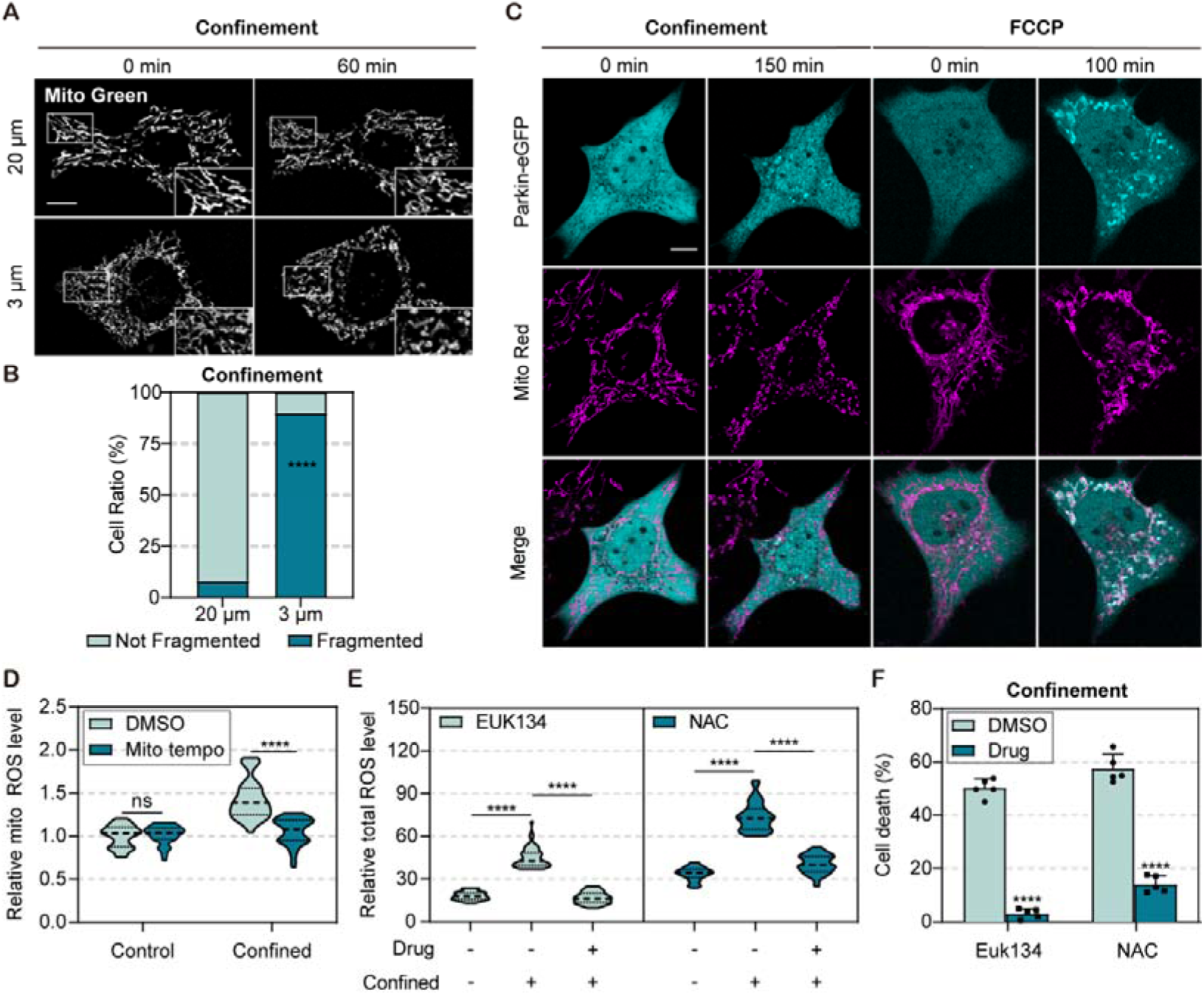
Confinement doesn’t trigger mitophagy and ROS scavengers inhibit cell death upon confinement. Related to Figure 3. (A) Representative time-lapse images of mitochondria (Mito Green) in HeLa cells upon 20 μm (top) or 3 μm (bottom) confinement. Insets are magnified images of areas indicated by white boxes. Scale bar = 10 μm. (B) Quantification of the percentage of cells with fragmented mitochondria in HeLa cells upon confinement for 60 min (*N* = 50 cells, *\*\*\*\*p* < 0.0001). (C) Representative live-cell images of the eGFP-Parkin expressing HeLa cells co-stained with Mito Red at 0 min and 180 min confinement. FCCP treatment (10 μM) is used as the positive control for mitophagy. Scale bar = 10 μm. (D) Relative mtROS level in HeLa cells treated with DMSO or MitoTEMPO (10 μM) upon confined and unconfined conditions. Data is normalized by the DMSO group without confinement (*N* = 40 cells, ns represents not significant, *\*\*\*\*p* < 0.0001). (E) Total ROS level in HeLa cells treated with DMSO or antioxidant drugs (EUK134, 100 μM; NAC, 500 μM) upon confined and unconfined conditions (*N* = 50 cells, *\*\*\*\*p* < 0.0001). (F) The percentage of cell death in HeLa cells treated with DMSO or antioxidant drugs upon confinement (*N* = 5 fields, *\*\*\*\*p* < 0.0001).

**Figure S4.**
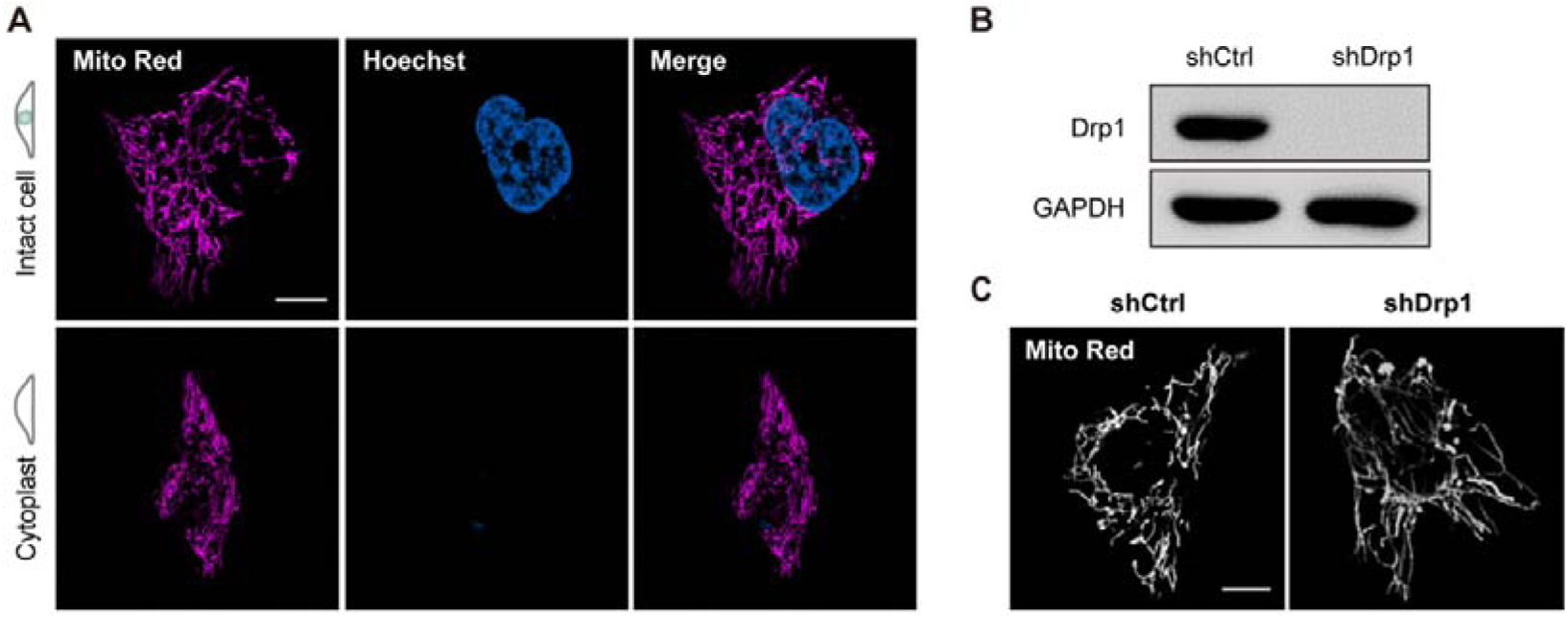
Enucleated cytoplasts preserve intact mitochondrial network. Related to Figure 4. (A) Representative live-cell images of mitochondria (Mito Red, magenta) in intact cells or enucleated cytoplasts. Scale bar = 10 μm. (B) Western blot shows the protein level of Drp1 in shCtrl and shDrp1 HeLa cells. GAPDH is used as the loading control. (C) Representative live-cell images of mitochondria (Mito Red) in shCtrl and shDrp1 HeLa cells. Scale bar = 10 μm.

**Figure S5.**
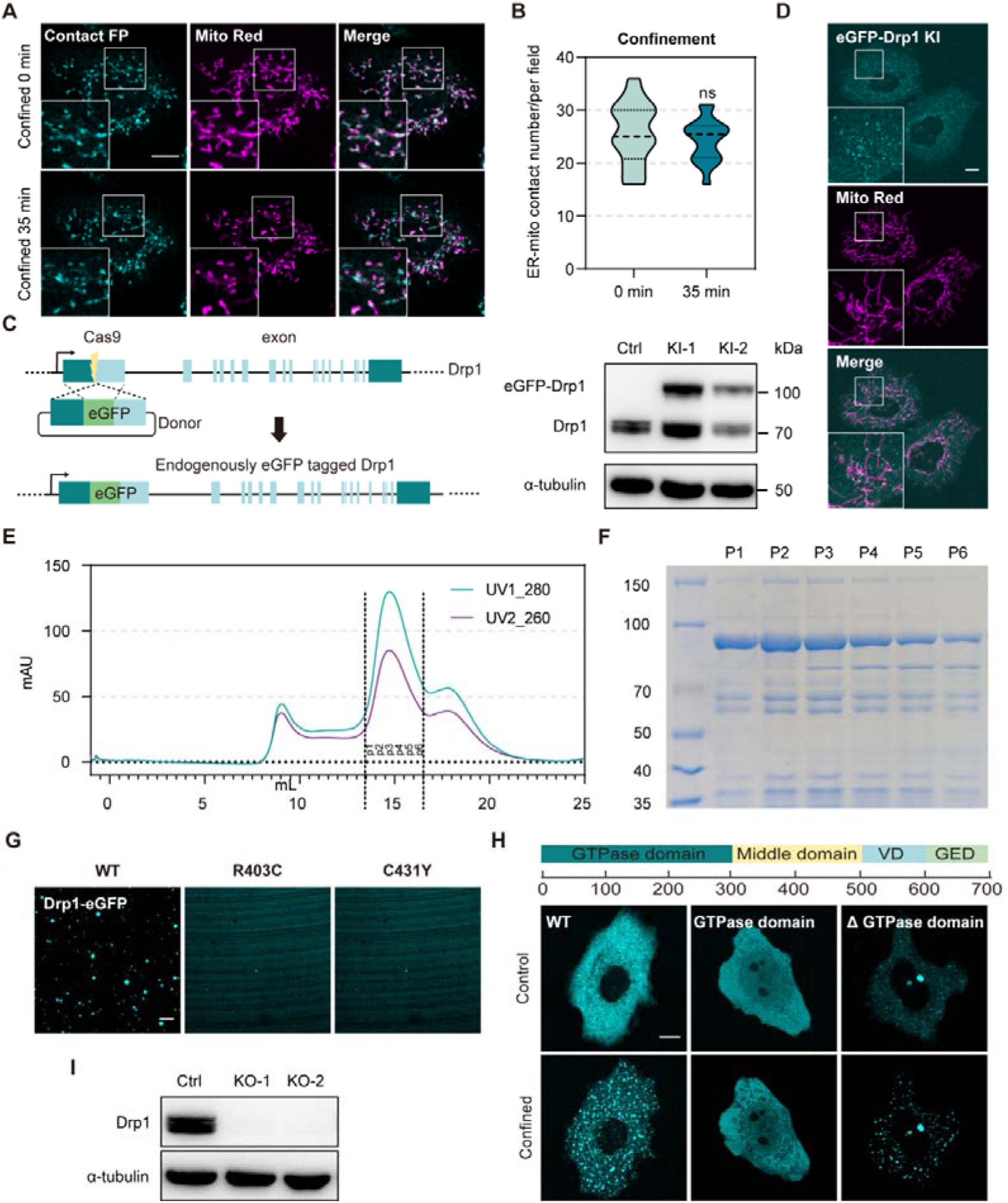
Drp1 point mutants in the middle domain inhibit its phase separation while the GTPase domain is dispensable for this property. Relate to Figure 5. (A) Representative time-lapse images of Mito-ER contacts and mitochondria (Mito Red) in HeLa cells co-transfected with pLVX-GA-Sec61β and pLVX-GB-MAVS upon confinement. Mito-ER contacts (cyan), Mitochondria (magenta). Insets are magnified images of areas indicated by white boxes. Scale bar = 10 μm. (B) Quantification of Mito-ER contacts number in HeLa cells at 0 min or 35 min confinement (*N* = 16 fields from 11 cells, ns represents no significant). (C) Left: schematic illustration of CRISPR-Cas9-mediated endogenous tagging of Drp1 gene with eGFP. A strategy consisting of sgRNA-guided Cas9 double-strand break (DSB) followed by homology-directed repair (HDR) is used. Right: Western blot shows the successful insertion of eGFP into the endogenous Drp1 locus. α-tubulin is used as the loading control. (D) Representative live-cell images of endogenous Drp1 and mitochondria (Mito Red) in HeLa cells. Scale bar = 10 μm. (E-F) Size-exclusion chromatography on Superose 6 (E) and Coomassie-blue-staining SDS-PAGE results (F) of Drp1-eGFP protein purified from *E.coli*. (G) Representative images of purified wild-type Drp1-eGFP (10 μM), R403C mutant (10 μM) or C431Y mutant (10 μM) diluted in 200 mM NaCl containing 8% dextran. Scare bar = 10 μm. (H) Top: Schematic representation of Drp1 domain structure. Bottom: Representative live-cell images of wild-type eGFP-Drp1, eGFP-GTPase domain, or eGFP-ΔGTPase domain expressing HeLa cells upon confined and unconfined conditions. Scare bar = 10 μm. (I) Western blot shows the protein level of Drp1 in wild-type and knock-out HeLa cells. α-tubulin is used as the loading control.

**Figure S6.**
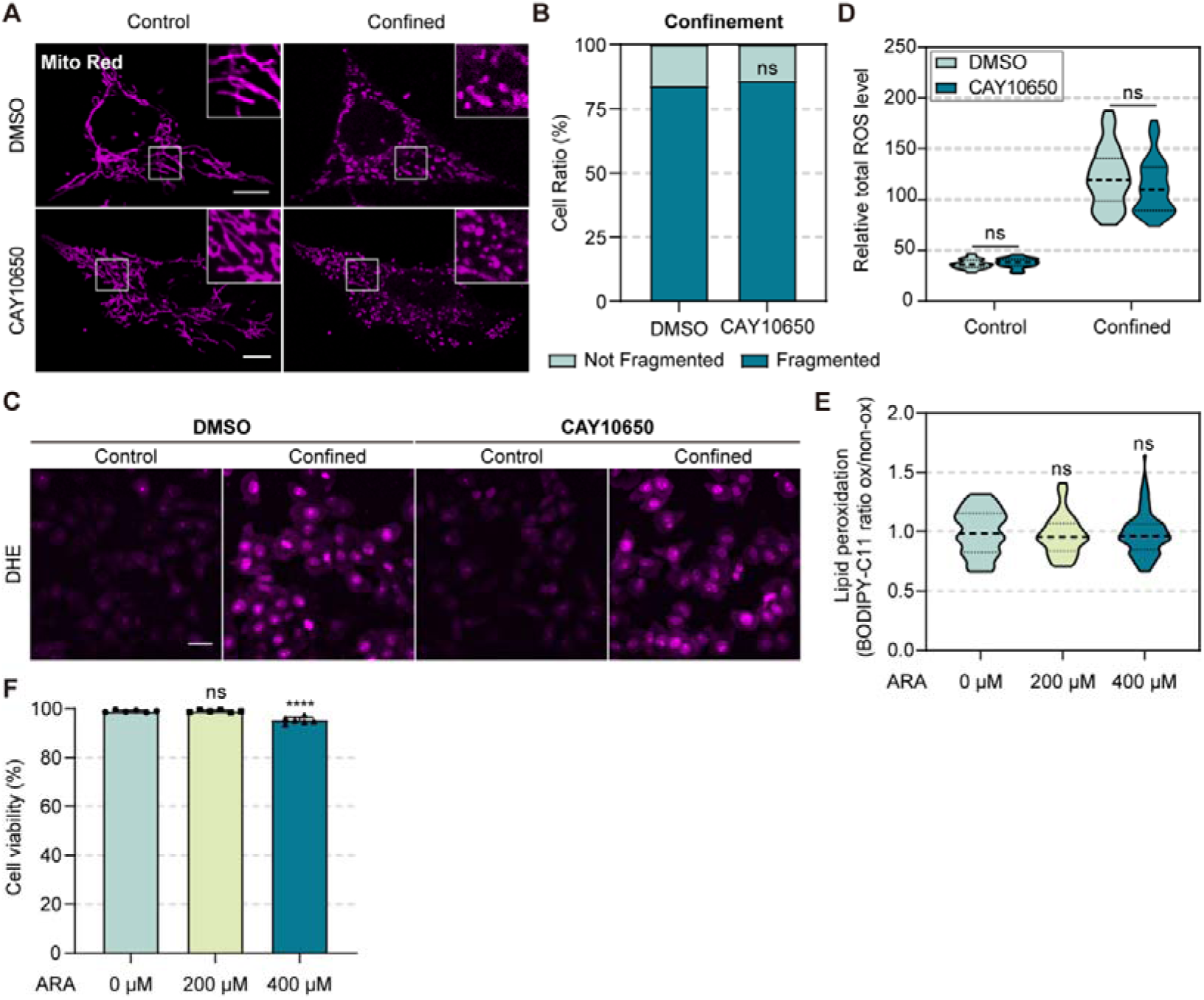
Ferroptosis cannot be induced solely by cPLA2 activation upon confinement. Related to Figure 6. (A) Representative live-cell images of mitochondria (Mito Red) in HeLa cells treated with DMSO or CAY10650 (150 nM). Insets are magnified images of areas indicated by white boxes. Scale bar = 10 μm. (B) Quantification of the percentage of cells with fragmented mitochondria in HeLa cells treated with DMSO or CAY10650 upon confinement (*N*_DMSO_ = 111 cells, *N*_CAY10650_ = 135 cells, ns represents not significant). (C) Representative live-cell images of DHE staining indicated total ROS level in HeLa cells treated with DMSO or CAY10650 upon confined and unconfined conditions. Scale bar = 40 μm. (D) Quantification of total ROS level in HeLa cells treated with DMSO or CAY10650 upon confinement (*N* = 40 cells, ns represents not significant). (E) Lipid ROS level in HeLa cells treated with ARA in gradient concentrations (0 μM, 200 μM and 400 μM, *N* = 55 cells, ns represents not significant). Cells in the 0 μM group are treated with DMSO for control. (F) Cell viability in HeLa cells treated with ARA in gradient concentrations (*N* = 6 fields, ns represents not significant, *\*\*\*\*p* < 0.0001). Cell viability (%) = (total cells-PI positive cells)/total cells * 100.

## Reference

1. Balzer, E.M. et al. Physical confinement alters tumor cell adhesion and migration phenotypes. The FASEB Journal 26, 4045–4056 (2012).

2. Uhler, C. & Shivashankar, G.V. Regulation of genome organization and gene expression by nuclear mechanotransduction. Nature Reviews Molecular Cell Biology 18, 717–727 (2017).

3. Liu, Y.J. et al. Confinement and low adhesion induce fast amoeboid migration of slow mesenchymal cells. Cell 160, 659–672 (2015).

4. Pagès, D.L. et al. Cell clusters adopt a collective amoeboid mode of migration in confined nonadhesive environments. Sci Adv 8 (2022).

5. Ekpenyong, A.E. et al. Mechanical deformation induces depolarization of neutrophils. Sci Adv 3 (2017).

6. Niethammer, P. Components and Mechanisms of Nuclear Mechanotransduction. Annual Review of Cell and Developmental Biology 37, 233–256 (2021).

7. Fletcher, D.A. & Mullins, D. Cell mechanics and the cytoskeleton. Nature 463, 485–492 (2010).

8. Humphrey, J.D., Dufresne, E.R. & Schwartz, M.A. Mechanotransduction and extracellular matrix homeostasis. Nature Reviews Molecular Cell Biology 15, 802–812 (2014).

9. Dupont, S. et al. Role of YAP/TAZ in mechanotransduction. Nature 474, 179–183 (2011).

10. Nava, M.M. et al. Heterochromatin-Driven Nuclear Softening Protects the Genome against Mechanical Stress-Induced Damage. Cell 181, 800–817.e822 (2020).

11. Lomakin, A.J. et al. The nucleus acts as a ruler tailoring cell responses to spatial constraints. Science 370, 310-+ (2020).

12. Venturini, V. et al. The nucleus measures shape changes for cellular proprioception to control dynamic cell behavior. Science 370, 311-+ (2020).

13. Denais, C.M. et al. Nuclear envelope rupture and repair during cancer cell migration. Science 352, 353–358 (2016).

14. Raab, M. ESCRT III repairs nuclear envelope ruptures during cell migration to limit DNA damage and cell death (vol 353, aah6167, 2016). Science 353, 1507–1507 (2016).

15. Nader, G.P.F. et al. Compromised nuclear envelope integrity drives TREX1-dependent DNA damage and tumor cell invasion. Cell 184, 5230–5246 e5222 (2021).

16. Nagata, S. & Tanaka, M. Programmed cell death and the immune system. Nature Reviews Immunology 17, 333–340 (2017).

17. Tang, R. et al. Ferroptosis, necroptosis, and pyroptosis in anticancer immunity. Journal of Hematology & Oncology 13 (2020).

18. Wang, S. et al. Mechanical overloading induces GPX4-regulated chondrocyte ferroptosis in osteoarthritis via Piezo1 channel facilitated calcium influx. Journal of Advanced Research 41, 63–75 (2022).

19. Le Berre, M., Aubertin, J. & Piel, M. Fine control of nuclear confinement identifies a threshold deformation leading to lamina rupture and induction of specific genes. Integr Biol (Camb) 4, 1406–1414 (2012).

20. Logue, J., Logue, J., Chadwick, R. & Waterman, C. A simple method for precisely controlling the confinement of cells in culture. Protocol Exchange (2018).

21. Wang, Y.P. et al. Chemotherapy drugs induce pyroptosis through caspase-3 cleavage of a gasdermin. Nature 547, 99-+ (2017).

22. Liu, X., Xia, S., Zhang, Z., Wu, H. & Lieberman, J. Channelling inflammation: gasdermins in physiology and disease. Nature Reviews Drug Discovery 20, 384–405 (2021).

23. Shi, J.J. et al. Cleavage of GSDMD by inflammatory caspases determines pyroptotic cell death. Nature 526, 660–665 (2015).

24. Zhang, Z. et al. Gasdermin E suppresses tumour growth by activating anti-tumour immunity. Nature 579, 415–420 (2020).

25. Sun, L. et al. Mixed Lineage Kinase Domain-like Protein Mediates Necrosis Signaling Downstream of RIP3 Kinase. Cell 148, 213–227 (2012).

26. Dixon, S.J. et al. Ferroptosis: An Iron-Dependent Form of Nonapoptotic Cell Death. Cell 149, 1060–1072 (2012).

27. Jiang, X., Stockwell, B.R. & Conrad, M. Ferroptosis: mechanisms, biology and role in disease. Nature Reviews Molecular Cell Biology 22, 266–282 (2021).

28. Hung, W.C. et al. Confinement Sensing and Signal Optimization via Piezo1/PKA and Myosin II Pathways. Cell Rep 15, 1430–1441 (2016).

29. Kalukula, Y., Stephens, A.D., Lammerding, J. & Gabriele, S. Mechanics and functional consequences of nuclear deformations. Nature Reviews Molecular Cell Biology 23, 583–602 (2022).

30. Enyedi, B., Jelcic, M. & Niethammer, P. The Cell Nucleus Serves as a Mechanotransducer of Tissue Damage-Induced Inflammation. Cell 165, 1160–1170 (2016).

31. Gaschler, M.M. & Stockwell, B.R. Lipid peroxidation in cell death. Biochem Bioph Res Co 482, 419–425 (2017).

32. Chao, C.C. et al. Metabolic Control of Astrocyte Pathogenic Activity via cPLA2-MAVS. Cell 179, 1483-+ (2019).

33. Graham, D.M. et al. Enucleated cells reveal differential roles of the nucleus in cell migration, polarity, and mechanotransduction. J Cell Biol 217, 895–914 (2018).

34. Efremov, Y.M., Kotova, S.L., Akovantseva, A.A. & Timashev, P.S. Nanomechanical properties of enucleated cells: contribution of the nucleus to the passive cell mechanics. J Nanobiotechnol 18 (2020).

35. Weindel, C.G. et al. Mitochondrial ROS promotes susceptibility to infection via gasdermin D-mediated necroptosis. Cell 185, 3214-+ (2022).

36. Lei, G., Zhuang, L. & Gan, B.Y. Targeting ferroptosis as a vulnerability in cancer. Nature Reviews Cancer 22, 381–396 (2022).

37. Doll, S. et al. ACSL4 dictates ferroptosis sensitivity by shaping cellular lipid composition. Nature Chemical Biology 13, 91–98 (2016).

38. Kraus, F., Roy, K., Pucadyil, T.J. & Ryan, M.T. Function and regulation of the divisome for mitochondrial fission. Nature 590, 57–66 (2021).

39. Smirnova, E., Griparic, L., Shurland, D.L. & van der Bliek, A.M. Dynamin-related protein Drp1 is required for mitochondrial division in mammalian cells. Molecular Biology of the Cell 12, 2245–2256 (2001).

40. Friedman, J.R. et al. ER Tubules Mark Sites of Mitochondrial Division. Science 334, 358–362 (2011).

41. Gregory E. Miner, S.Y.S., Wendy K. Showalter, Christina M. So, J.V.R., Alex E. Powers, Maria Clara Zanellati, & Chih-Hsuan Hsu, M.F.M., and Sarah Cohen Contact-FP: A dimerization-dependent fluorescent protein toolkit for visualizing membrane contact site dynamics. Contact 7: 1–12 (2024).

42. Shin, Y. et al. Spatiotemporal Control of Intracellular Phase Transitions Using Light-Activated optoDroplets. Cell 168, 159-+ (2017).

43. Bauer, B.L., Rochon, K., Liu, J.C., Ramachandran, R. & Mears, J.A. Disease-associated mutations in Drp1 have fundamentally different effects on the mitochondrial fission machinery. Hum Mol Genet 32, 1975–1987 (2023).

44. Korobova, F., Ramabhadran, V. & Higgs, H.N. An Actin-Dependent Step in Mitochondrial Fission Mediated by the ER-Associated Formin INF2. Science 339, 464–467 (2013).

45. Wong, Y.C., Ysselstein, D. & Krainc, D. Mitochondria–lysosome contacts regulate mitochondrial fission via RAB7 GTP hydrolysis. Nature 554, 382–386 (2018).

46. Rowland, A.A. & Voeltz, G.K. Endoplasmic reticulum–mitochondria contacts: function of the junction. Nature Reviews Molecular Cell Biology 13, 607–615 (2012).

47. Alisafaei, F., Jokhun, D.S., Shivashankar, G.V. & Shenoy, V.B. Regulation of nuclear architecture, mechanics, and nucleocytoplasmic shuttling of epigenetic factors by cell geometric constraints. P Natl Acad Sci USA 116, 13200–13209 (2019).

48. Quirós, P.M., Mottis, A. & Auwerx, J. Mitonuclear communication in homeostasis and stress. Nature Reviews Molecular Cell Biology 17, 213–226 (2016).

49. Mohammadi, H. & Sahai, E. Mechanisms and impact of altered tumour mechanics. Nature Cell Biology 20, 766–774 (2018).

50. Berg, S. et al. ilastik: interactive machine learning for (bio)image analysis. Nature Methods 16, 1226–1232 (2019).

