## Supplementary Movies for "Mechanical confinement induces ferroptosis through mitochondrial dysfunction"

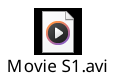

Movie S1. Cells blew blebs from the plasma membrane and gradually lysed upon confinement, related to Figure 1. Time-lapse movie showing the morphology dynamic of HeLa cells at 3 μm confinement. Nuclei: Hoechst, blue; Propidium iodide: magenta. White arrow: plasma membrane blebs. Time interval: 3 min, total time: 9 hours. Scale bar: 20 μm.

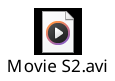

Movie S2. Ferroptosis inhibitors abrogate confinement-induced cell death, related to Figure 1. Time-lapse movie showing the morphology dynamic of HeLa cells treated with DMSO or ferroptosis inhibitors at 3 μm confinement, including DFOM, Ferrostatin-1, and Liproxstatin-1. Nuclei: Hoechst, blue; Propidium iodide: magenta. Time interval: 3 min, total time: 12 hours. Scale bar: 20 μm.

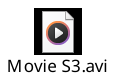

Movie S3. cPLA2 translocates to mitochondria from cytoplasm upon confinement, related to Figure 2. Time-lapse movie of cPLA2-eGFP expressing HeLa cells co-stained with Mito Red at 3 μm confinement. Mito Red: magenta; cPLA2-eGFP: cyan. Time interval: 2 min, total time: 2 hours. Scale bar: 10 μm.

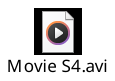

Movie S4. Mitochondria fragment upon confinement, related to Figure 3. Time-lapse movie of HeLa cells transfected with Tom20-mCherry at 3 μm confinement. Time interval: 2 min, total time: 90 min. Scale bar: 5 μm.

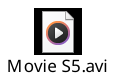

Movie S5. Drp1 is recruited and forms puncta at mitochondrial fission sites upon confinement, related to Figure 5. Time-lapse movie of eGFP-Drp1 knock-in HeLa cells co-stained with Mito Red at 3 μm confinement. Mito Red: magenta; eGFP-Drp1: cyan. White arrowhead: the formation of bright Drp1 puncta at mitochondrial sites which subsequently underwent fission. Time interval: 20 s, total time: 40 min. Scale bar: 5 μm.
